# Comprehensive 3D mapping reveals distinct spatial gradients of SST, PV, and TH interneurons across the mouse caudoputamen

**DOI:** 10.64898/2026.01.25.701604

**Authors:** Jonibek M. Muhsinov, Evan A. Iliakis, Wenxin Tu, Alexandra N. Ramirez, Michael Muniak, Andrzej Wasilczuk, M. Felicia Davatolhagh, Alexander Proekt, Tianyi Mao, Marc V. Fuccillo

## Abstract

In addition to spatially organized excitatory forebrain inputs along its mediolateral, dorsoventral, and anteroposterior axes, the striatum relies on cellular diversity to subserve its myriad processing functions. Distinct striatal GABAergic interneuron subtypes, including somatostatin (SST), parvalbumin (PV), and tyrosine hydroxylase (TH) interneurons likely subserve complementary computational roles. However, a detailed understanding of how these microcircuit components are distributed across striatal territories remains lacking. To address this gap, we generated a comprehensive three-dimensional atlas of SST, PV, and TH interneurons across the mouse caudoputamen using genetic labeling, brain-wide imaging, and voxel-wise quantification. We found that SST and TH interneurons were relatively enriched in the ventral caudoputamen, whereas PV interneurons were enriched dorsally. In addition, PV and TH interneurons exhibited opposing anteroposterior distribution patterns, with PV interneurons enriched posteriorly and TH interneurons showing a marked decline in density towards the striatal tail. Consequently, while the three interneuron subtypes displayed comparable densities in the functionally defined lateral striatum and anterior ventromedial striatum, PV interneurons predominated in the dorsomedial striatum and tail of striatum. We did not observe major sex differences. Together, these findings reinforce the view that the striatum is not a monolithic structure: in addition to structured excitatory inputs, inhibitory microcircuits themselves are differentially distributed across striatal territories, providing region-specific constraints on circuit computation. By integrating interneuron organization into existing anatomical frameworks, this atlas provides a foundation for linking striatal anatomy to function across behavioral domains.

## Introduction

The striatum is the main input nucleus of the basal ganglia. Striatal principal neurons—the spiny projection neurons (SPNs)—and sparse cholinergic and heterogeneous GABAergic interneuron subtypes process distributed forebrain projection inputs to support learning, reward processing, movement, and cognition (Choi et al., 2018; Cox & Witten, 2019; Graybiel & Grafton, 2015; Klug et al., 2018; Martel & Galvan, 2022; Wall et al., 2013). Importantly, the striatum is not a monolith. Anatomic tracing studies reveal a striking spatial organization of cortical and thalamic inputs, largely supporting the notion that more dorsolateral striatal regions subserve sensorimotor functions, while more ventromedial regions subserve more limbic functions (Hintiryan et al., 2016; Hunnicutt et al., 2016). These findings are bolstered by decades of functional data suggesting that the dorsolateral striatum, dorsomedial striatum, and ventral striatum respectively support sensorimotor, associative, and limbic roles (Alexander et al., 1991; Cox & Witten, 2019). There is also evidence indicating that the anteroposterior axis represents an additional dimension of striatal organization, with anterior and posterior territories subserving distinct functions during behavior (Cifuentes et al., 2025; Corbit & Janak, 2010; Mestres-Missé et al., 2012; Yin et al., 2005). Notably, cortical projections do not follow a simple nearest-neighbor mapping along the anteroposterior axis. Instead, distinct projection-defined neuronal subpopulations within a single cortical region preferentially innervate anterior versus posterior striatal territories and exhibit dissociable task-related representations (Beckstead, 1979; Choi et al., 2023; Kruttner et al., 2022; Selemon & Goldman-Rakic, 1985), underscoring the AP axis as a meaningful anatomical reference frame with functional relevance (Cifuentes *et al*., 2025; Shiflett et al., 2010; Wang et al., 2018; Yin *et al*., 2005).

In addition to spatial organization of inputs, the striatum relies on cellular diversity (Tepper et al., 2018; Tepper et al., 2010) to subserve its myriad processing functions. While the functional properties of SPNs, cholinergic interneurons, and midbrain dopaminergic afferents are comparatively well understood (Apicella, 2017; Cox & Witten, 2019; Lee & Sabatini, 2025), recent work reveals surprising roles for sparse, local circuit GABAergic interneurons in a range of striatal processes. Dendritic-targeting somatostatin (SST) interneurons modulate corticostriatal signaling and learning-related network reorganization (Fino et al., 2018; Holly et al., 2019; Holly et al., 2021; Rotariu et al., 2025; Straub et al., 2016). Soma-targeting parvalbumin (PV) interneurons exert powerful control over spike timing and coordinated ensemble activity (Duhne et al., 2021; Duhne et al., 2024; Duhne et al., 2025; Gritton et al., 2019; Martiros et al., 2018; O’Hare et al., 2017; Owen et al., 2018). A third interneuron class unique to the striatum, Tac2+ nondopaminergic tyrosine hydroxylase (TH) interneurons (Corrigan et al., 2025; Ibanez-Sandoval et al., 2010; Xenias et al., 2015), remains less well characterized, but has been shown to engage local striatal networks and influence behavior (Appings et al., 2024; Ibanez-Sandoval et al., 2010; Kaminer et al., 2019), notably via inhibitory interactions with SST interneurons (Assous et al., 2017).

While found throughout the striatum, the distribution of these interneuron subtypes is not uniform and not thoroughly described. Previous studies have identified regional biases and planar gradients in interneuron density (Bernácer et al., 2007; 2012; Lecumberri et al., 2017; López-González del Rey et al., 2022; Luk & Sadikot, 2001; Monteiro et al., 2018; Muñoz-Manchado et al., 2018; Unal et al., 2011; Van Zandt et al., 2024; Wu & Parent, 2000), yet a unified framework of their organization across the full three-dimensional extent of the striatum (as exists for cholinergic interneurons; see Carrasco et al., 2022; Hobel et al., 2026; Matamales et al., 2016) remains lacking. Because distinct interneuron classes regulate different aspects of local circuit function, including dendritic integration and network synchronization, their relative abundance and spatial arrangement could dictate the types of computations made by each striatal territory. A detailed understanding of how these microcircuit components are distributed across striatal territories therefore provides critical anatomical constraints for interpreting circuit-level function and for contextualizing experimental findings across studies.

To address this gap, we generated a comprehensive three-dimensional atlas of SST, PV, and TH interneurons across the mouse caudoputamen using genetic labeling, brain-wide imaging, and voxel-wise quantification. This analysis revealed distinct spatial principles governing interneuron organization, including axis-specific gradients and regionally defined subtype compositions. This resource provides an anatomical framework for interpreting and hypothesizing about interneuron function across striatal territories.

## Materials and Methods

### Animals

All procedures and experiments were conducted in accordance with the National Institutes of Health Guidelines for the Use of Animals and approved by the University of Pennsylvania Institutional Animal Care and Use Committee (Protocol: 805643). All mice used in this study were aged 42-43 days at the time of perfusion and were on the C57BL/6J genetic background.

To label and visualize the spatial distribution of somatostatin (SST), parvalbumin (PV), and tyrosine hydroxylase (TH) interneurons, male mice homozygous for the Cre-reporter line Ai14D (Gt(ROSA)26Sor(tm14(CAG-tdTomato)Hze)/J; Jax 007914) were crossed with female mice expressing either: (1) SST-ires-Cre (Sst(tm2.1(cre)/Zjh)/J; Jax 013044), (2) PV-2a-Cre (Pvalb(tm1.1(cre)Aibs)/J; Jax 012358), or (3) BAC-transgenic TH-Cre (Tg(Th-cre)Fl12Gsat/Mmucd); MMRRC 037415-UCD). This breeding scheme was used to mitigate risk the of germline recombination in PV-2a-Cre x Ai14D crosses (Kobayashi & Hensch, 2013).

SST-ires-Cre; Ai14D heterozygous, PV-2a-Cre; Ai14D heterozygous, and TH-Cre; Ai14D heterozygous offspring were analyzed. Both male and female mice were included. The sensitivity and specificity of SST-ires-Cre and PV-2a-Cre lines for labeling dorsal striatal SST+ and PV+ interneurons have been validated previously (Choi et al., 2018). The BAC-transgenic TH-Cre line has been used previously to study striatal TH+ interneurons (Kaminer et al., 2019; Xenias et al., 2015).

### Tissue Preparation for Anatomic Mapping

Mice were deeply anesthetized via intraperitoneal injection of pentobarbital sodium (150 μL; Sagent, NDC # 25021-676-20). Following loss of the toe-pinch response, mice were transcardially perfused with 15 mL Formalin (10% phosphate-buffered; Fisher Scientific SF100-4) mixed with heparin (50 μL of 1000 USP units/mL; Meitheal 71288-402-11). Brains were extracted and post-fixed in Formalin overnight (12-24 hours).

Following fixation, brains were sectioned coronally at 50μm using a Vibratome (5100 mz; Campden Instruments) in phosphate-buffered saline (PBS). Free-floating sections were serially mounted and coverslipped with Fluoromount-G (Southern Biotech 0100-01) containing DAPI (0.6 μM; Fisher Scientific D1306). Sections were imaged and stitched using a Leica DM6 epifluorescence microscope at 10x magnification.

### Immunohistochemistry

Mice were transcardially perfused with 1x phosphate-buffered saline (PBS), followed by 4% paraformaldehyde (PFA). Brains were extracted and post-fixed in 4% PFA for 2 hours. Tissue was sectioned coronally at 50μm using a vibratome (Vibratome, Model 1000 Plus) in PBS.

Free-floating sections were permeabilized with 0.2% Triton X-100 and blocked for 1 hour in 3% normal goat serum (NGS) in PBS. Sections were incubated overnight with primary antibody (rat monoclonal anti-somatostatin, 1:500, Millipore, #MAB354) diluted in PBS containing 1% NGS and 0.2% Triton X-100. After washing, sections were incubated with secondary antibody for 2 hours (goat anti-rat IgG (H+L), Alexa Fluor 555 conjugate, 1:500, Invitrogen A21434), then mounted and imaged using an Olympus BX63 epifluorescence microscope at 10x magnification.

### 3D Reconstruction, Atlas Registration, and Cell Detection

Following tissue preparation and imaging, exported TIFF files were converted to MBF NeuroInfo-compatible JPEG 2000 format using MBF MicroFile+ and imported into MBF NeuroInfo for all subsequent reconstruction, atlas registration, and cell detection analyses. General data processing workflows using MBF NeuroInfo have been described previously (Eastwood et al., 2018).

Serial sections were assembled and reconstructed using the Serial Section Assembler tool. Individual sections were initially outlined automatically based on image contrast and edge detection, then manually refined as needed. Sections were ordered and aligned to generate a three-dimensional reconstruction, with additional manual refinement performed post-alignment to ensure consistent orientation and section spacing. Three-dimensional reconstructions were saved as JPEG 2000 image stacks (JPX).

Reconstructed brains were manually registered to the Allen Mouse Brain Common Coordinate Framework (CCFv3) using the Register Sections tool in MBF NeuroInfo (version 2024.1.3). Section angle was determined using the following anatomical landmarks: (1) the section at which the corpus callosum begins to cross the midline, (2) the section in which the rostral anterior commissure appears as three separate components, and (3) the first section in which the fasciculus retroflexus appears as a compact, rounded fiber bundle. The remaining sections were registered automatically and manually refined to maintain consistent alignment and spacing across the entire brain.

Neurons expressing tdTomato were detected using MBF NeuroInfo’s Cell Detection Workflow tool. For SST-Cre; Ai14D brains, image intensity ranges were adjusted (0-500 a.u.) to minimize detection of tdTomato-positive but SST-non-immunoreactive cell clusters, which likely reflect developmentally restricted SST expression (see Supplemental Figure 1). SST-Cre tdTomato-positive cells were detected using a diameter range of 10-15.5 μm and a fixed sensitivity of 11,500 (see Eastwood et al., 2018 for detailed description of this procedure).

For PV-Cre; Ai14D brains, three separate cell detection workflows were performed per brain to minimize detection of fibers of passage as PV interneurons. These workflows were defined based on reproducible qualitative changes in PV-positive non-somatic process morphology observed across serial sections, including changes in process caliber, density, continuity, and orientation relative to section plane. Specifically, PV-positive non-somatic processes transition from relatively sparse, fine-caliber, and discontinuous profiles to denser, thicker processes that increasingly form continuous fiber-like structures, and ultimately to long, thin processes that run predominantly parallel to the section plane. Boundaries between workflows were determined by visual inspection of these morphological transitions (see Figure 1C). PV-Cre tdTomato-positive cells were detected using a diameter range of 7.5-15 μm, with sensitivities optimized separately for each workflow. For TH-Cre; Ai14D brains, cells were detected using a diameter range of 7-24 μm, with sensitivities optimized on a per-brain basis.

**Figure 1.**
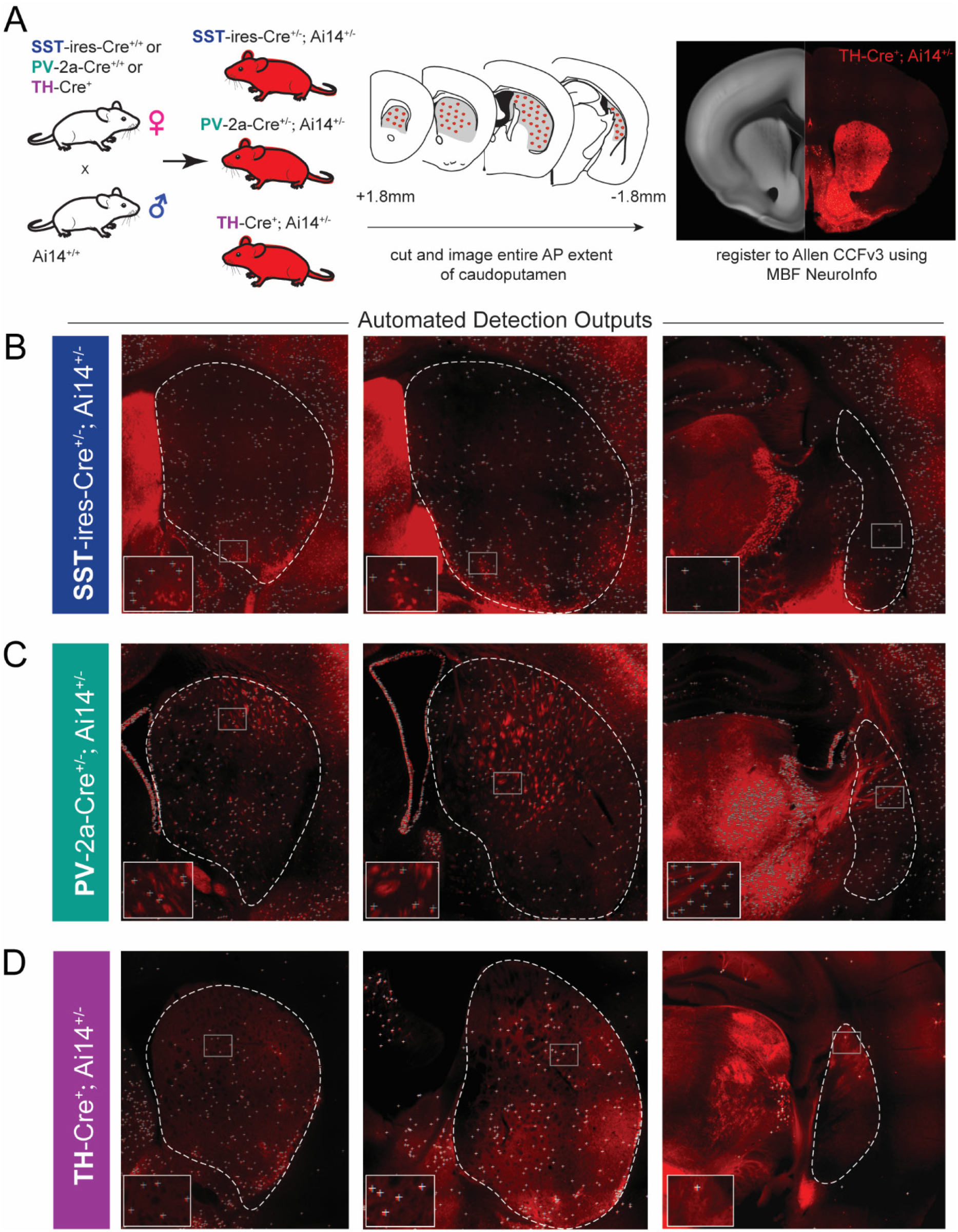
Workflow for genetic identification and brain-wide mapping of SST, PV, and TH interneurons in the mouse caudoputamen. (A) Breeding strategy and overview of tissue preparation, imaging, and atlas registration pipeline. (Left) Mice expressing Cre recombinase in SST, PV, or TH interneurons were crossed to Ai14D reporter mice, and coronal sections spanning the full anteroposterior extent of the caudoputamen (middle) were imaged and registered to The Allen Mouse Brain Common Coordinate Framework (CCFv3) using MBF NeuroInfo (right). (B-D) Representative epifluorescence images illustrating detection of SST+ (B), PV+ (C), and TH+ (D) cells. Caudoputamen extent is delineated by dotted white lines. The lower left inset is a magnification of boxes within the striatum (‘+’ mark detected cells). Detection parameters were optimized separately for each interneuron subtype to minimize false-positive detection. For SST interneurons, settings were adjusted to reduce detection of tdTomato-positive but SST-immunonegative cell clusters (see Supplemental Fig 1). For PV interneurons, detection parameters were optimized to minimize inclusion of non-somatic PV+ axons and dendrites.

Cell detection outputs were exported for further analysis as CSV files containing per-cell anteroposterior (AP), mediolateral (ML), and dorsoventral (DV) coordinates in Allen CCFv3 reference space. Individual CSV files can be shared upon request.

### Anatomical Data Analysis

All subsequent anatomical data analysis was performed in MATLAB using custom scripts, available upon request. Cell coordinate outputs from MBF NeuroInfo were filtered to include only detections classified as caudoputamen. All left hemisphere cell detections were reflected to the right hemisphere prior to analysis.

To estimate cell density, hemisphere-level observations were resampled with replacement (1,000 bootstrapped samples) and binned into cubic voxels (150 × 150 × 150 µm). Voxel-wise densities (cells/mm^3^) were computed separately for each cell type. Voxels were masked using Allen CCFv3 caudoputamen and ventricle masks (structures 672 and 73 respectively, available at https://download.alleninstitute.org/informatics-archive/current-release/mouse_ccf/annotation/ccf_2017/structure_masks/structure_masks_50/) and excluded if their Chebyshev distance was less than 50 µm from the caudoputamen boundary or less than 175 µm from the ventricle. The ventricle mask was especially important for exclusion of non-neuronal PV-positive ependymal cells from PV interneuron density estimates. Density maps were smoothed using a Gaussian filter (σ = 0.5 voxel).

Density values were subsequently pooled across all cell types to generate 15 quantile-based density thresholds, which were visualized as heatmaps and voxel-wise predominance heatmaps in Figure 2. Collapsed one-dimensional density profiles were generated for Figure 3, with the standard error of bootstrapped density estimates calculated and overlaid.

**Figure 2.**
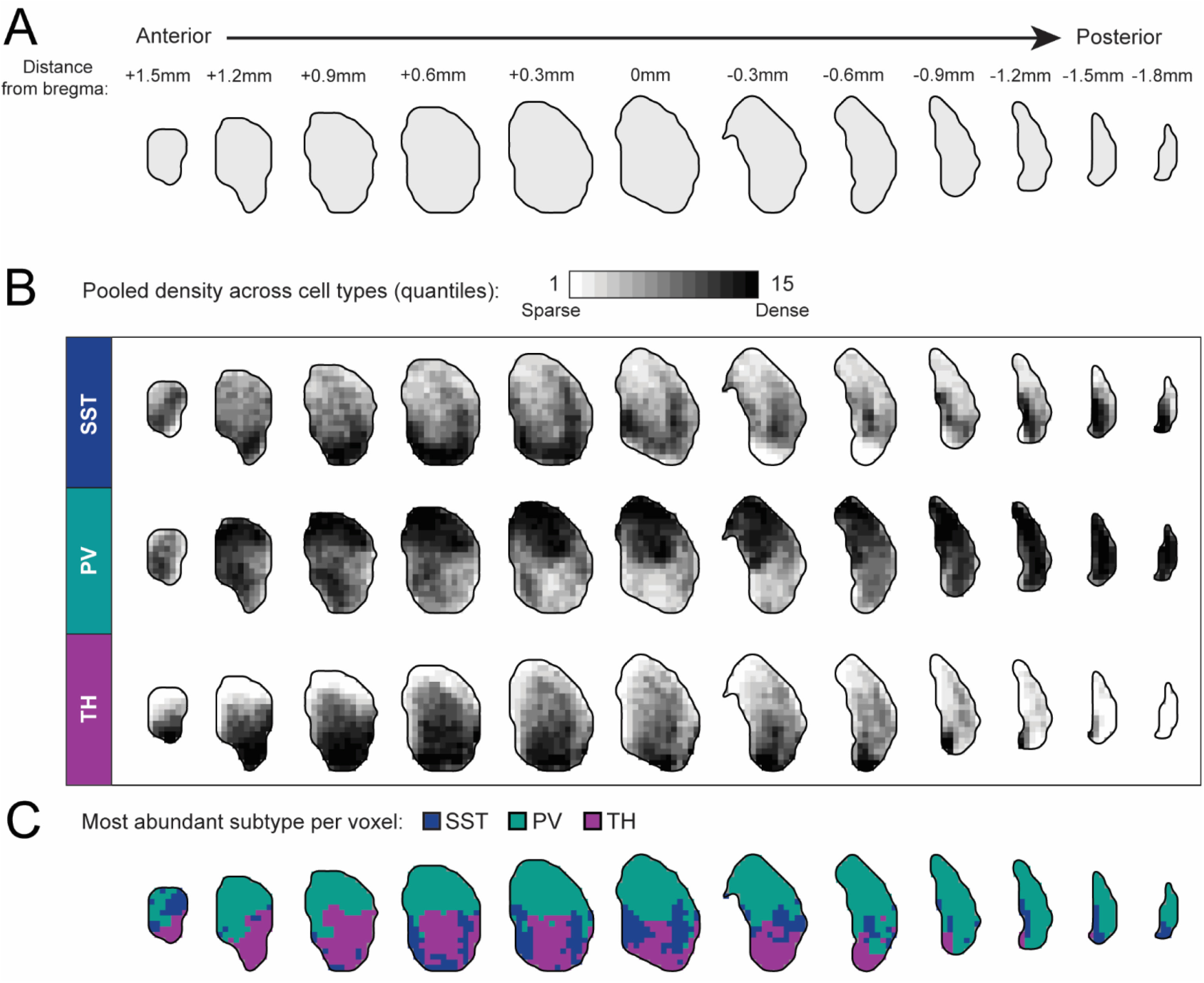
Comprehensive three-dimensional atlas of SST, PV, and TH interneuron distribution across the mouse caudoputamen. (A) Schematic illustrating the anteroposterior extent of the caudoputamen sampled in coronal planes shown at 300 μm intervals, with distance to bregma (mm) indicated above each plane. (B) Voxel-wise density maps showing the spatial distribution of SST (*N*(hemispheres) = 12), PV (*N*(hemispheres) = 11), and TH (*N*(hemispheres) = 13) interneurons across the mouse caudoputamen. The caudoputamen was partitioned into 150-μm voxels, and interneuron densities were computed for each voxel. Density values were subsequently pooled across all cell types to generate 15 quantile-based density thresholds, visualized from sparse to dense. Heatmaps disaggregated by sex are shown in Supplemental Figure 2. (C) Voxel-wise predominance maps, in which each voxel was assigned to the interneuron subtype exhibiting the highest density. Voxels are color-coded to indicate the most abundant interneuron subtype (blue, SST; green, PV; purple, TH). This predominance map provides a qualitative summary of relative enrichment and does not imply exclusivity of interneuron subtypes within individual voxels. Predominance maps disaggregated by sex are shown in Supplemental Figure 3.

**Figure 3.**
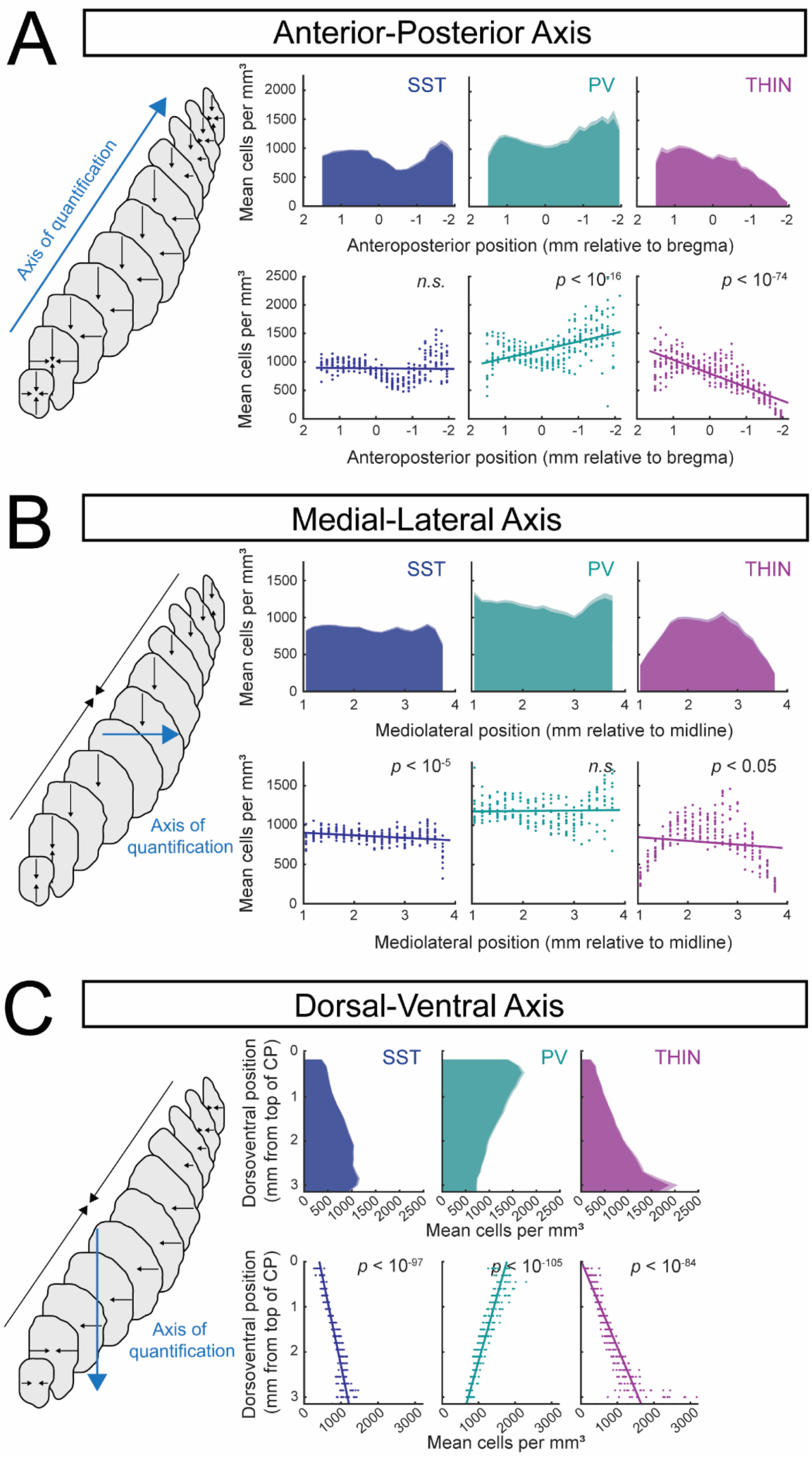
Large-scale spatial gradients of striatal interneuron subtypes across anatomical axes. Voxel-wise interneuron density estimates were quantified along the anterior-posterior (A), medial-lateral (B), and dorsal-ventral (C) axes of the mouse caudoputamen. Left schematizes the axis of quantification (blue line) and the axes of compression (black lines). Top rows depict bootstrapped mean density for SST (N(hemispheres) = 12), PV (N(hemispheres) = 11), and TH (N(hemispheres) = 13) across 150-μm planes (1,000 bootstraps per plane, error bars indicate s.e. of the bootstrapped mean). Plots disaggregated by sex are shown in Supplemental Figure 4. Bottom rows show per-hemisphere raw density values for each plane, with overlaid linear mixed-effects model fits. Linear mixed-effects models (Density ∼ Coordinate + (1 | Hemisphere_ID)) were used to assess overall directional bias along each anatomical axis. Model slopes represent global directional bias (cells/mm^3^ per mm along each anatomical axis) and are not intended to capture non-monotonic structure. Data disaggregated by sex are shown in Supplemental Figure 5. Anterior-posterior axis. SST: β = 4.9 ± 11.1, p = 0.66 (N = 12 hemispheres); PV: β = −146.2 ± 16.1, p = 3.0 × 10^-17^ (N = 11 hemispheres); TH: β = 239.6 ± 9.7, p = 3.9 × 10^-75^ (N = 13 hemispheres). Medial-lateral axis. SST: β = −32.4 ± 7.0, p = 6.7 × 10^-6^ (N = 12 hemispheres); PV: β = 6.5 ± 14.0, p = 0.65 (N = 11 hemispheres); TH: β = −47.5 ± 21.6, p = 0.029 (N = 13 hemispheres). Dorsal-ventral axis. SST: β = 253.9 ± 7.3, p = 3.9 × 10^-98^ (N = 12 hemispheres); PV: β = −350.7 ± 8.7, p = 4.0 × 10^-106^ (N = 11 hemispheres); TH: β = 519.7 ± 17.9, p = 5.6 × 10^-85^ (N = 13 hemispheres).

Cell coordinates were additionally assigned to one of four striatal subregions (see Figure 4A) based on previously defined clusters from Hunnicutt et al. (2016). These subregions were transformed into Allen CCFv3 space and masked using the Allen CCFv3 caudoputamen mask to generate subregion volumes. Per-hemisphere densities (cells/mm^3^) were calculated for each cell type and subregion and visualized as boxplots (Figure 4B).

**Figure 4.**
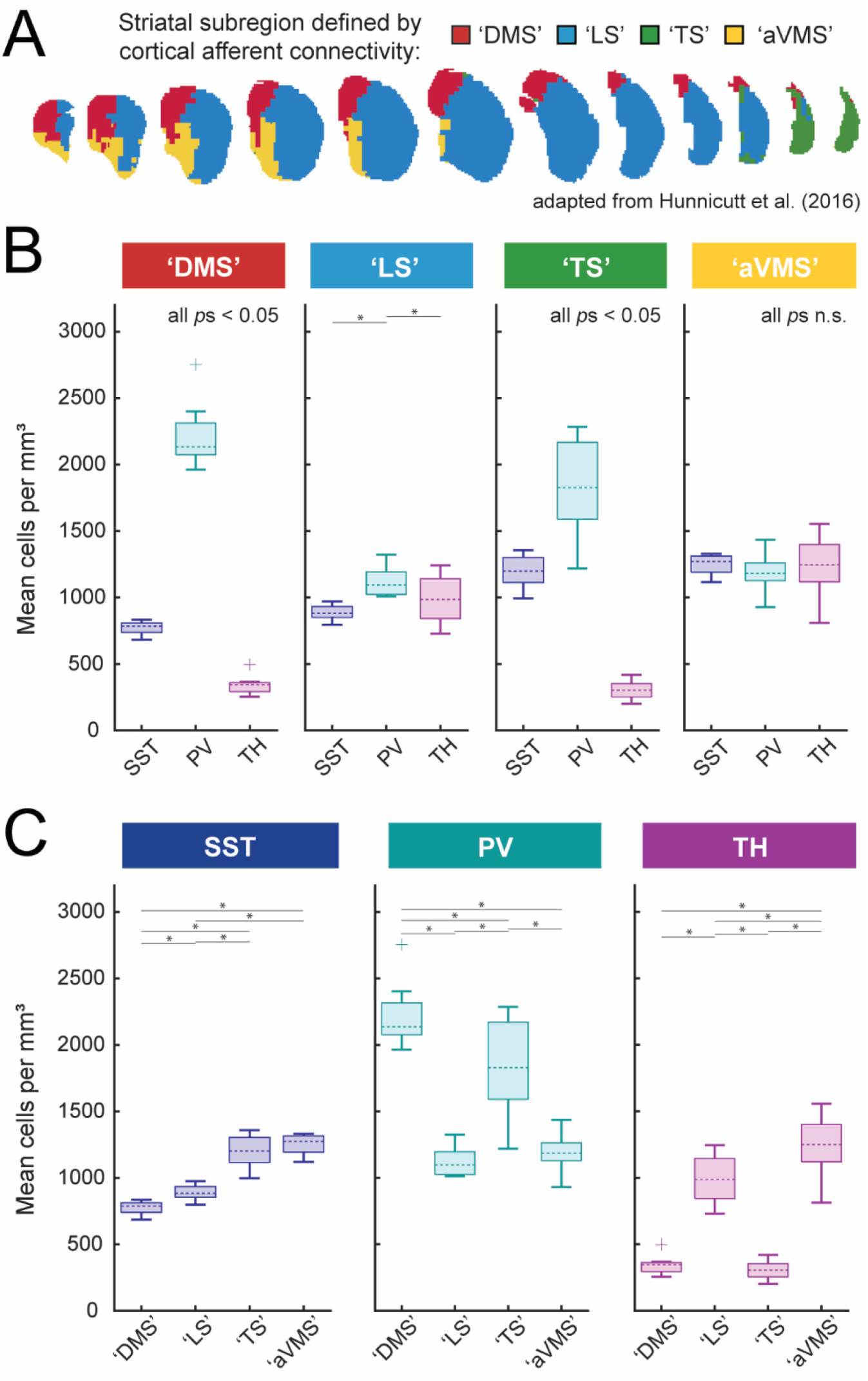
Subregional organization of striatal SST, PV, and TH interneurons. (A) Schematic adapted from Hunnicutt et al. (2016) illustrating a four-cluster anatomic parcellation of the caudoputamen based on cortical input patterns. Accumbal territories were excluded using a caudoputamen mask. We adopt the following labels: Cluster 5 (red), dorsomedial striatum (DMS); Cluster 7 (blue), lateral striatum (LS); Cluster 12 (green), tail of striatum (TS); Cluster 15 (gold), anterior ventromedial striatum (aVMS). (B-C) Per-hemisphere subregional density distributions (boxplots; center line, median; box, interquartile range; whiskers, non-outlier extrema; points, outliers) are shown organized by subregion (B) or by interneuron subtype (C). Mixed-effects ANOVAs (linear mixed-effects models with random intercept for hemisphere; (1 | Hemisphere_ID)) were used to test (i) subtype differences within each subregion (Density ∼ Subtype + (1 | Hemisphere_ID)) and (ii) subregional differences within each subtype (Density ∼ Subregion + (1 | Hemisphere_ID)). Sidak-corrected significant pairwise comparisons (*p* < 0.05) are marked with asterisks. Omnibus subtype effects within subregion (panel B): DMS: *p* = 2.1 × 10^-22^; LS: *p* = 3.9 × 10^-4^; TS: *p* = 2.7 × 10^-14^; aVMS: *p* = 0.32 (n.s.). Omnibus subtype effects within subtype (panel C): SST: *p* = 2.1 × 10^-15^; PV: *p* = 6.5 × 10^-13^; TH: *p* = 6.4 × 10^-22^.

### Statistical Methods

To quantify the large-scale spatial organization of interneuron density along major striatal axes, voxel-wise densities were projected onto the corresponding anatomical coordinates and analyzed using linear mixed-effects models. This approach was chosen to provide a parsimonious, directionally interpretable summary of axis-level bias, rather than to model fine-scale spatial structure.

First-pass models included anatomical coordinate as the sole fixed effects predictor, with hemisphere identity included as a random intercept to account for repeated voxel measurements across hemispheres:

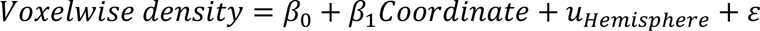

Follow-up models to investigate sex differences included anatomical coordinate, sex, and their interaction as fixed effects, with hemisphere identity included as a random intercept to account for repeated voxel measurements across hemispheres:

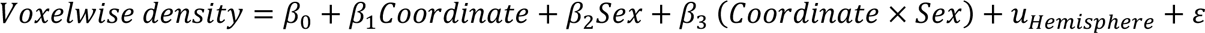

Models were fit using the *fitlme* function in MATLAB (formula first-pass: “Density ∼ Coordinate * Sex + (1 | Hemisphere_ID)”; formula follow-up: “Density ∼ Coordinate * Sex + (1 | Hemisphere_ID)”).

A similar approach was used to examine differences in interneuron composition across cortical projection-defined striatal territories, both within subregions (cell-type composition) and across subregions (regional enrichment). In these analyses, interneuron density was modeled as a function of either interneuron subtype (for within-subregion comparisons), or subregion (for within-subtype comparisons), together with sex and their interactions in follow-up models, with hemisphere identity included as a random intercept.

## Results

To characterize the distribution of somatostatin (SST), parvalbumin (PV), and tyrosine hydroxylase (TH) interneurons throughout the mouse caudoputamen, we generated a three-dimensional atlas using genetic labeling, brain-wide imaging, and voxel-wise quantification. To this end, mice homozygous for SST-Cre, PV-Cre, or TH-Cre were crossed to the Cre-dependent reporter line Ai14D (**Fig 1A**), yielding offspring in which targeted interneuron populations expressed the red fluorophore tdTomato.

At post-natal day 42-43, mice were sacrificed, and coronal sections containing the caudoputamen were imaged and registered to the Allen Common Coordinate Framework v3 (CCFv3) using MBF NeuroInfo (full detains in Methods). The built-in Cell Detection Workflow was used to identify tdTomato-positive somata and map their spatial coordinates into CCFv3 space (Fig 1B-D).

Although the SST-ires-Cre line has previously been validated for sensitivity and specificity to dorsal striatal SST interneurons (Choi et al., 2018; Holly et al., 2019), we observed discrete clusters of dense tdTomato-positive somata largely in ventrolateral regions of our SST-Cre; Ai14D mice that lacked immunoreactivity for SST (Supplemental Fig 1). These clusters likely reflect developmentally restricted SST or non-specific expression. Regardless, to ensure conservative quantification, detection parameters were optimized to minimize the contribution of these clusters to our final cell counts (see Fig 1B for an example of excluded signal, and Supplemental Fig 1 for a detailed description of the exclusion procedure).

In our PV-2a-Cre; Ai14D crosses, we additionally observed tdTomato-positive fiber-like structures lacking identifiable somata, with morphology and density that varied across the anteroposterior axis of the striatum. These structures were distinct from the compact, round profiles characteristic of neuronal cell bodies and likely reflect labeling of axonal processes from PV-expressing populations outside the striatum. To prevent contamination of voxel-wise cell density estimates, detection parameters were optimized to selectively exclude non-somatic signal while preserving bona-fide tdTomato-positive cell bodies (see Fig 1C and Supplemental Fig 1).

### Voxel-wise three-dimensional map of GABAergic interneuron distributions

To generate a three-dimensional map of striatal SST, PV, and TH interneuron distributions, we partitioned the caudoputamen into 150-μm voxels and computed the voxel-wise interneuron densities across the striatum (Fig 2).

SST interneurons exhibited a non-uniform distribution, with relative enrichment in the ventral portion of caudoputamen, and particularly within the anterior dorsomedial striatum (DMS), as well as additional regions of elevated density along the lateral striatum. SST interneurons were also prominent within the medial of the striatal tail (Fig 2B, top). In contrast, PV interneurons were distributed broadly throughout the dorsal striatum across anteroposterior levels, with comparatively lower density in ventral regions. Notably, PV density remained high in posterior striatum and extended throughout the striatal tail (Fig 2B, middle). TH interneurons showed the most circumscribed spatial profile, with very sparse representation in dorsal- and medial-most aspects of the striatum, and near absence from the striatal tail. Instead, the highest TH interneuron densities were observed in ventral and more anterior planes of the striatum (Fig 2B, bottom).

In anterior sections, SST and TH interneurons displayed partially overlapping ventral enrichment patterns that contrasted with more dorsally biased distribution of PV interneurons. This organization progressively diminished in posterior planes as TH interneuron density declined (Fig 2B).

These spatial relationships are summarized using a voxel-wise predominance map (Fig 2C) in which each voxel is assigned to the interneuron subtype exhibiting the highest normalized density. This representation highlights the predominance of PV interneurons in the dorsal striatum, relative enrichment of SST interneurons in ventral and posterior regions, and preferential representation of TH interneurons in anterior and ventral striatum. Importantly, the predominance map is intended as a qualitative summary of relative enrichment and does not imply exclusivity of interneuron subtypes within individual voxels. Atlas and predominance maps disaggregated by sex are provided in Supplemental Figures 2, 3.

### Interneuron organization along major striatal axes

To quantify large-scale spatial organization of striatal interneuron subtypes, we examined how SST, PV, and TH interneuron densities vary along the anteroposterior (AP), mediolateral (ML), and dorsoventral (DV) axes of the striatum (Fig 3). To address whether interneurons were preferentially enriched towards one extreme of an anatomical axis relative to the other, voxel-wise densities were projected onto the corresponding anatomical coordinates (blue ‘axis of quantification’) and analyzed using linear mixed-effects models. This approach was chosen to provide a parsimonious, directionally interpretable summary of axis-level bias, rather than to model fine-scale spatial structure. Because several distributions exhibited clear non-monotonic features, model slopes were interpreted as measures of global directional tendency rather than complete descriptions of spatial patterning.

### Anteroposterior axis

Along the anteroposterior axis, both SST and PV interneurons exhibited non-uniform distributions characterized by a local reduction in density at intermediate striatal levels, and a marked increase toward posterior, tail-associated regions (Fig 3A). This posterior enrichment was particularly prominent for PV interneurons. Interestingly, the increase in SST cell density towards the tail was restricted to more medial parts of this structure. Consistent with this pattern, linear mixed-effects modeling revealed a significant overall increase in PV density along the anteroposterior axis (β = - 146.2, *p* = 3.0 × 10^-17^, *N*(hemispheres) = 11, *N*(sections) = 297), whereas SST interneurons did not exhibit a significant monotonic change across the axis (β = 4.9, *p* = 0.66, *N*(hemispheres) = 12, *N*(sections) = 324).

In contrast, TH interneurons displayed a pronounced decrease in density toward posterior striatal levels (Fig 3A; β = 239.6, *p* = 3.9 × 10^-75^, *N*(hemispheres) = 13, *N*(sections) = 351), consistent with atlas-level distributions shown in Fig 2B. This posterior decline was significantly steeper in males than females (β_female = 212.9, β_male = 270.75, sex x position β = 57.85; *p* = 0.00278). Sex-disaggregated data are provided in Supplemental Figures 4, 5.

### Mediolateral axis

Along the mediolateral axis, SST interneurons exhibited relatively uniform density, with a small but statistically significant negative slope (Fig 3B; β = −32.4, *p* = 6.7×10^-6^, *N*(hemispheres) = 12, *N*(sections) = 264), indicating subtle lateral depletion rather than a strong spatial gradient.

PV interneurons showed a clearly non-monotonic mediolateral distribution, with elevated density only in the lateral-most planes (Fig 3B). In line with this nonlinear structure, linear mixed-effects modeling did not detect a significant overall slope across the mediolateral axis (β = 6.5, *p* = 0.65, *N*(hemispheres) = 11, *N*(sections) = 242). Notably, males and females exhibited divergent trends across the mediolateral axis (β_female = −61.34, β_male = 62.96, sex x position interaction β = 124.3, *p* = 5.66 × 10^-6^), with sex-disaggregated data distributions shown in Supplemental Figures 4, 5.

TH interneurons displayed reduced density at both the most medial- and lateral-most planes, with higher density in intermediate mediolateral positions (Fig 3B), consistent with atlas-level observations (Fig 2B). This nonlinear pattern yielded a small but statistically significant negative slope in linear mixed-effects modeling (β = −47.5, *p* = 0.029, *N*(hemispheres) = 11, *N*(sections) = 242), likely reflecting reduced TH density in tail regions.

### Dorsoventral axis

Along the dorsoventral axis, both SST and TH interneurons exhibited robust ventral enrichment. TH interneuron density increased steeply toward ventral striatum (β = 519.7, *p* = 5.6 × 10^-85^, *N*(hemispheres) = 13, *N*(sections) = 299), while SST interneurons showed a more moderate but highly significant ventral increase (β = 253.9, *p* = 3.9 × 10^-98^, *N*(hemispheres) = 12, *N*(sections) = 276). The dorsoventral TH gradient was significantly more pronounced in males than females (β_female = 441.6, β_male = 610.8, sex x position interaction β = 169.2, *p* = 1.54 × 10^-6^); sex-disaggregated data are shown in Supplemental Figures 4, 5.

In contrast, PV interneurons exhibited a strong decrease in density along the dorsoventral axis (β = - 350.7, *p* = 4.0 × 10^-106^, *N*(hemispheres) = 11, *N*(sections)= 253), indicating preferential dorsal localization relative to SST and TH interneurons.

### Subregion-level organization of striatal interneuron subtypes

Another approach to quantify large-scale spatial gradients was with regard to cortical afferent-defined territories (as previously examined in Hunnicutt *et al*., 2016). Striatal voxels were assigned to sub-clusters ‘dorsomedial striatum’ (DMS), ‘lateral striatum’ (LS), ‘tail of striatum’ (TS), and ‘anterior ventromedial striatum’ (aVMS) clusters based on dominant cortical afferent identity derived from Allen Institute connectivity mapping studies (Fig 4A). Linear mixed-effects models were used to assess differences in interneuron density both within defined subregions (cell-type composition; Fig 4B) and across subregions (regional enrichment; Fig 4C).

### Within-subregion interneuron composition

Within dorsal striatal subregions, interneuron composition was broadly similar. In both the dorsomedial striatum (DMS) and tail striatum (TS), PV interneurons were the most abundant subtype, SST interneurons were present at intermediate densities, and TH interneurons were sparsest (Fig 4B), consistent with the dorsal bias observed in axis-based analyses.

In the lateral striatum (LS), subtype distinctions were less pronounced. PV enrichment was reduced, and SST and TH interneurons were present at more comparable densities. A significant cell type x sex interaction (*F*(2,30) = 4.8226, *p* = 0.015), was driven by a greater density of TH interneurons in males (mean density = 1087.6 cells/mm^3^; 95% CI [887.6, 1287.5]) than in females (mean density = 904.4 cells/mm^3^; 95% CI [803.4, 1005.4]), as shown in Supplemental Figures 6, 7. In males, TH and PV interneurons were similarly abundant (*p*_Sidak_ = 1), and each exceeded SST interneuron density (*p*s_Sidak_ < 0.03), whereas females followed the general population pattern with PV predominance.

In the anterior ventromedial striatum (aVMS), interneuron densities were comparatively balanced across SST, PV, and TH subtypes. A significant cell type x sex interaction (*F*(2,30) = 4.9432, *p* = 0.014) reflected greater TH interneuron density in males (mean density = 1364.2 cells/mm^3^; 95% CI [1239.3, 1489.2]) than in females (m(density) = 1163.1 cells/mm^3^; 95% CI [935.4, 1390.7]), as shown in Supplemental Figures 6, 7. In males, TH interneurons were significantly more abundant than PV interneurons (*p*_Sidak_ < 0.005), whereas no subtype-specific differences were detected in females.

### Across-subregion comparisons

Across corticostriatal territories, interneuron subtypes exhibited distinct regional biases (Fig 4C). SST interneurons were most abundant in aVMS and TS and were sparsest in DMS. PV interneurons reached their highest densities in DMS and TS and were less abundant in LS and aVMS. In contrast, TH interneurons displayed a complementary distribution relative to PV interneurons, with lower density in DMS and TS and greater representation in LS and aVMS.

Together, these results indicate that continuous spatial gradients in interneuron density resolve into distinct subtype compositions across cortical afferent-defined striatal territories. Thus, large-scale anatomical organization of interneuron populations is reflected not only along continuous coordinate axes, but also in discrete functional architecture defined by cortical input domains.

## Discussion

In this study, we generated a three-dimensional atlas of somatostatin (SST), parvalbumin (PV), and tyrosine hydroxylase (TH) interneurons in the dorsal striatum of mice using genetic labeling, brain-wide imaging, and voxel-wise analyses. We quantified gradients of distribution using linear-mixed effects models and compared interneuron densities across cortical afferent-defined striatal subregions, finding striking differences in the distribution of the three interneurons subtypes (Fig 2). Most notably, we found that SST and TH interneurons were relatively enriched in the ventral striatum, whereas PV interneurons were enriched dorsally (Fig 3C). PV and TH interneurons also had opposite anteroposterior distribution patterns, with PV interneurons enriched posteriorly while TH interneurons showed a significant decline in density posterior to bregma and were nearly absent from the striatal tail (Fig 3A). Thus, whereas the three interneuron subtypes had comparable densities in the functionally defined ‘lateral striatum’ (LS) and ‘anterior ventromedial striatum’ (aVMS), PV interneurons predominated in the ‘dorsomedial striatum’ (DMS) and ‘tail of striatum’ (TS; Fig 4). We did not observe major sex differences in our dataset.

In several respects, our findings align with previously published literature. We recapitulate a dorsal-to-ventral gradient of decreasing PV interneuron density (López-González del Rey et al., 2022; Luk & Sadikot, 2001; Tepper et al., 2010), as well as an anteroposterior gradient of increasing PV interneuron density, consistent with observations in macaques (López-González del Rey et al., 2022) but differing from earlier findings in rats (Wu & Parent, 2000). Importantly, the latter study did not include the striatal tail; in our dataset, a pronounced increase in PV interneuron density in this region was the primary contributor to the observed anteroposterior gradient, likely accounting for the discrepancy between studies. We additionally observe a dorsal-to-ventral gradient of increasing density in TH interneurons, as previously described in macaques (López-González del Rey et al., 2022). To our knowledge, no studies have quantitatively assessed distribution gradients of SST interneurons across the caudoputamen. However, one recent study found that SST-immunoreactive cells are more abundant in the nucleus accumbens than in the caudoputamen (Van Zandt et al., 2024), hinting at a dorsal-to-ventral gradient of increasing density as observed in our analysis.

Some aspects of our findings diverge from prior anatomic studies, however. With our recombinase-based genetic strategy, we do not recapitulate the medial-to-lateral gradient of increasing density of PV interneurons that has been reported in immunohistochemical (IHC) studies in mice (Monteiro et al., 2018), rats (not Garas et al., 2016; but Gerfen, 1985; Wu & Parent, 2000), squirrel monkeys (Wu & Parent, 2000), macaques (López-González del Rey et al., 2022), and humans (Bernácer et al., 2012). Because our approach provides a binarized readout of PV expression, rather than quantitative information about expression levels, one possible explanation for this discrepancy is that IHC studies may be sensitive to gradients in PV expression rather than differences in absolute interneuron number (Monteiro et al., 2018). This interpretation is supported by single-cell RNA-sequencing analyses, demonstrating that *Pvalb* is highly co-expressed with *Pthlh*, and that neurons expressing these markers constitute a shared PV/PTHLH transcriptional class (Muñoz-Manchado et al., 2018). Within this class, *Pthlh* transcript levels decrease along the mediolateral axis, contrasting with an increase in *Pvalb* transcript density along the same dimension. Moreover, genetic labeling in PV-Cre mice captured interneurons that were *Pthlh+* but *Pvalb-* by in situ hybridization, suggesting that IHC or ISH for *Pvalb* alone might fail to identify even genetically labeled interneurons in this class. One explanation for the identified PV gradient is that *Pvalb* expression levels scale with electrophysiological demands associated with rapid excitatory drive. Such an interpretation is borne out by Patch-Seq in this same study (Muñoz-Manchado et al., 2018), which finds that higher *Pvalb* transcript levels were correlated with lower action potential half-width and higher maximal firing rates in the PV/PTHLH class. Together, these findings raise the possibility that mediolateral gradients observed in PV IHC studies reflect variation in PV expression levels within a distributed interneuron class, rather than differences in absolute cell density.

Because distinct interneuron classes regulate different aspects of local circuit function, including dendritic integration and network synchronization, their relative abundance and spatial arrangement could dictate the types of computations that predominate in each striatal territory. Accordingly, a detailed understanding of how these microcircuit components are distributed across striatal territories therefore provides important anatomical constraints for interpreting circuit-level function and for contextualizing experimental findings across studies. For example, dendritic-targeting somatostatin (SST) interneurons modulate corticostriatal signaling and learning-related network reorganization (Fino et al., 2018; Holly et al., 2019; Holly et al., 2021; Rotariu et al., 2025; Straub et al., 2016), while soma-targeting parvalbumin (PV) interneurons exert powerful control over spike timing and coordinated ensemble activity (Duhne et al., 2021; Duhne et al., 2025; Gritton et al., 2019; Martiros et al., 2018; O’Hare et al., 2017; Owen et al., 2018). The relative enrichment of PV interneurons in the dorsal striatum may therefore reflect greater computational demands for precise regulation of ensemble dynamics in sensorimotor processing. Conversely, the increased representation of SST interneurons in ventral regions may be consistent with heightened demands on regulation of dendritic excitability and synaptic plasticity in territories canonically associated with reward-processing and learning. However, it is important to note that these constraints are not deterministic. While interneuron distributions may inform hypotheses about the relative computational capacities of different striatal subregions, they are not prescriptive. For example, even in regions where particular interneuron subtypes are comparatively sparse, their density remains non-null, indicating that regional differences are quantitative rather than categorical. Furthermore, specific interneuron subtypes may exhibit synaptic connectivity at a distance from their cell bodies (Straub et al., 2016). Ultimately, in vivo recordings and manipulations will be necessary to discern functional contributions of interneuron subtypes (e.g., Duhne et al., 2025; Holly et al., 2019; Lee et al., 2017; Owen et al., 2018).

Our study has several limitations that should be considered. Firstly, the use of recombinase-based genetic strategies carries the inherent risk of labeling neurons with developmentally restricted gene expression that are not bona fide members of the interneuron class of interest. For example, in our SST-Cre; Ai14D mice, we observed dense clusters of tdTomato+ cells that were not SST-immunoreactive and whose distribution and morphology were inconsistent with prior descriptions of striatal SST interneurons (Supplemental Figure 1). In addition, genetic labeling results in expression within extrinsic axonal or dendritic processes, as well as occasional non-neuronal cell types, which can complicate automated cell detection and quantification. This issue was most pronounced in our PV-Cre; Ai14D mice, where periventricular ependymal cells and dense PV+ axons necessitated masking and more rigorous detection methods to isolate PV+ somata. Although we implemented conservative exclusion criteria (Supplemental Figure 1), it remains possible that these factors introduced bias into density estimates.

While our TH-Cre; Ai14D line did not exhibit such issues, the high density of TH+ dopamine neuron axons made it impossible for us to immunohistochemically validate the sensitivity and specificity of this line to TH interneurons. However, the TH-Cre line used here has been widely employed in prior studies of striatal TH interneurons (Assous et al., 2017; Kaminer et al., 2019; Xenias et al., 2015), and the distribution patterns we observed—including ventral enrichment and relative sparing of the tail and dorsal- and medial-most aspects of striatum—closely mirror *Th* mRNA expression patterns in the Allen Mouse Brain Atlas (Lein et al., 2006).

Additional methodologic limitations include reconstruction of three-dimensional distributions patterns from sectioned tissue, lack of intersectional genetic or immunohistochemical validation, and reliance on SST, PV, and TH as primary markers of interneuron identity. Future work could leverage tissue clearing and light sheet imaging approaches (e.g., CLARITY, iDISCO), combined with whole-brain immunohistochemistry or in situ hybridization for complementary markers, including nNOS, NPY, and CHODL for SST interneurons; PTHLH for PV interneurons, TAC2 for TH interneurons, and GAD67 for all, to provide more robust classification of interneuron subtypes. Finally, our analysis did not address the compartmental organization of interneurons within striosomal and matrix domains, another important principle of striatal organization (Brimblecombe & Cragg, 2017; Graybiel & Matsushima, 2023). For example, TH interneurons have been reported to preferentially localize to striosomes in ventral striatum (Unal et al., 2011), a feature we were unable to evaluate here. Future studies integrating interneuron mapping with markers such as the μ-opioid receptor may help resolve how interneuron distributions interact with striatal compartmentalization.

Together, these findings reinforce the view that the striatum is not a monolithic structure but is instead organized along multiple spatial dimensions that shape how information is locally processed. In addition to structured excitatory inputs, inhibitory microcircuits themselves are differentially distributed across striatal territories, providing region-specific constraints on circuit computation. By integrating these microcircuit features into existing anatomical frameworks, this atlas provides a foundation for linking striatal anatomy to function across behavioural domains.

## Conflict of Interest

The authors declare that the research was conducted in the absence of any commercial or financial relationships that could be construed as a potential conflict of interest.

## Funding

This project was supported by NIMH Grant F30MH136699 to Evan Iliakis and NIMH Grants R01MH118369 and RF1MH138591 to Marc Fuccillo.

## Acknowledgments

We would like to thank Alessandro Jean-Louis for outstanding technical assistance, and Maxime Assous and Fulva Shah for TH-Cre transgenic mice. We also thank MBF Bioscience for providing technical support with the NeuroInfo interface. We would also like to acknowledge the use of OpenAI’s ChatGPT (versions 4o, 5, 5.1, 5.2) as an auxiliary tool for code optimization and editorial refinement.

## Data Availability Statement

The datasets generated for this study are available upon request.

## Supplemental Figures

**Supplemental Figure 1.**
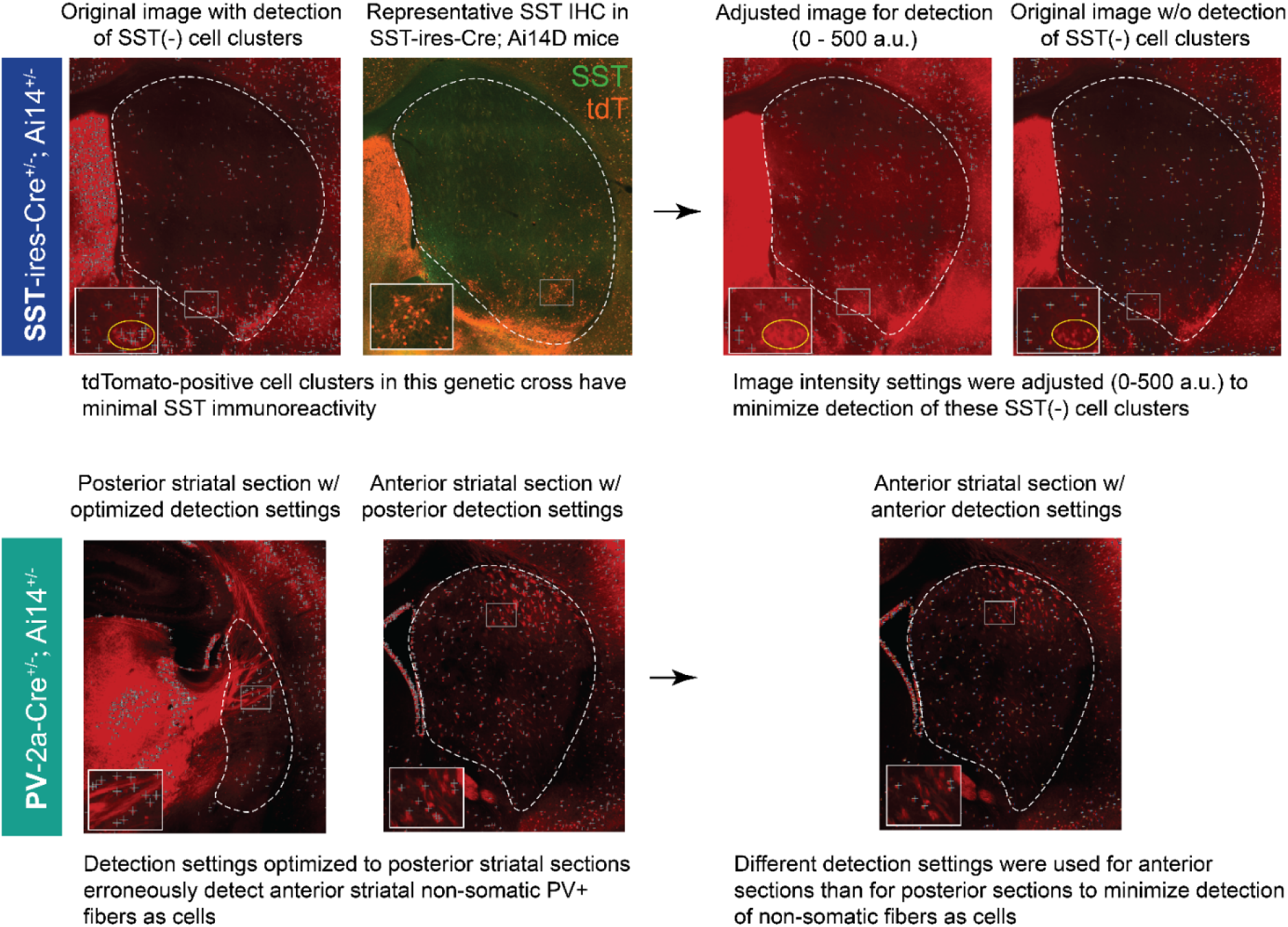
Optimization of detection parameters for SST and PV interneurons. Row 1 (SST interneurons): In SST-ires-Cre; Ai14D mice, tdTomato-positive cell clusters were observed that lacked detectable SST immunoreactivity. To restrict automated detection to SST-immunopositive somata, image intensity display ranges were adjusted (0-500 a.u.) such that SST-negative clusters were excluded from detection. This approach substantially reduced their contributions to final SST interneuron counts. Row 2 (PV interneurons): Non-somatic PV+ fiber bundles exhibit distinct morphology and vary along the anteroposterior axis of the striatum. Detection parameters optimized for posterior sections minimized inclusion of these fibers but resulted in over-detection when applied to anterior planes. To avoid systemic inclusion of non-somatic PV-positive structures, separate detection sensitivity settings were applied for anterior, middle, and posterior sections, ensuring preferential detection of PV-positive somata across the full anteroposterior extent of the striatum.

**Supplemental Figure 2.**
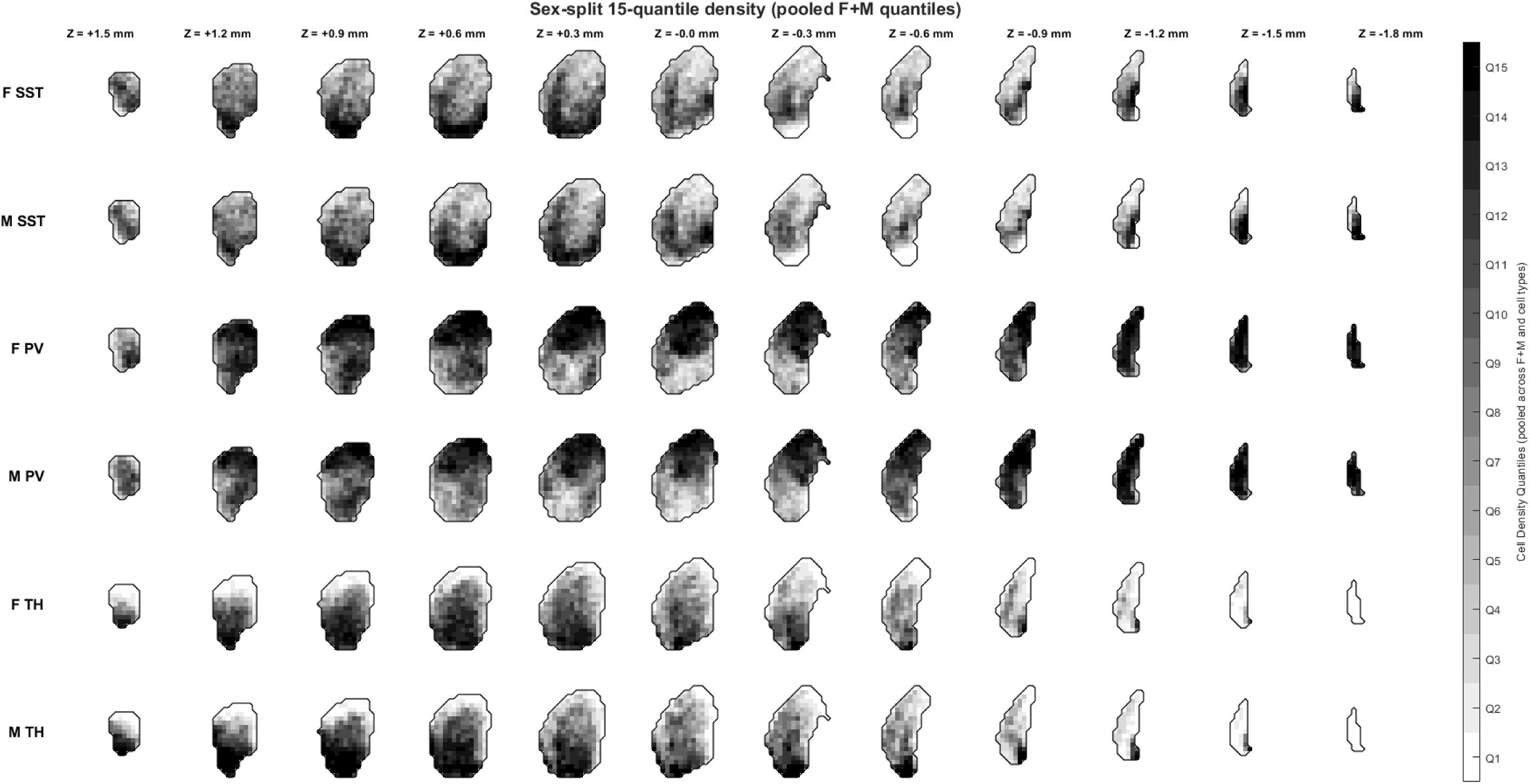
Comprehensive three-dimensional atlas of SST, PV, and TH interneuron distribution across the mouse caudoputamen, stratified by sex. Voxel-wise density maps showing the spatial distribution of SST (*N*(hemispheres) = 12; 6 female, 6 male), PV (*N*(hemispheres) = 11; 5 female, 6 male), and TH (*N*(hemispheres) = 13; 7 female, 6 male) interneurons across the mouse caudoputamen. The caudoputamen was partitioned into 150-μm voxels, and interneuron densities were computed for each voxel. For visualization purposes, density values were pooled across cell types and sexes to generate 15 quantile-based density thresholds, displayed from sparse to dense. This common quantile scale enables qualitative comparison of large-scale spatial organization across interneuron subtypes and sexes.

**Supplemental Figure 3.**
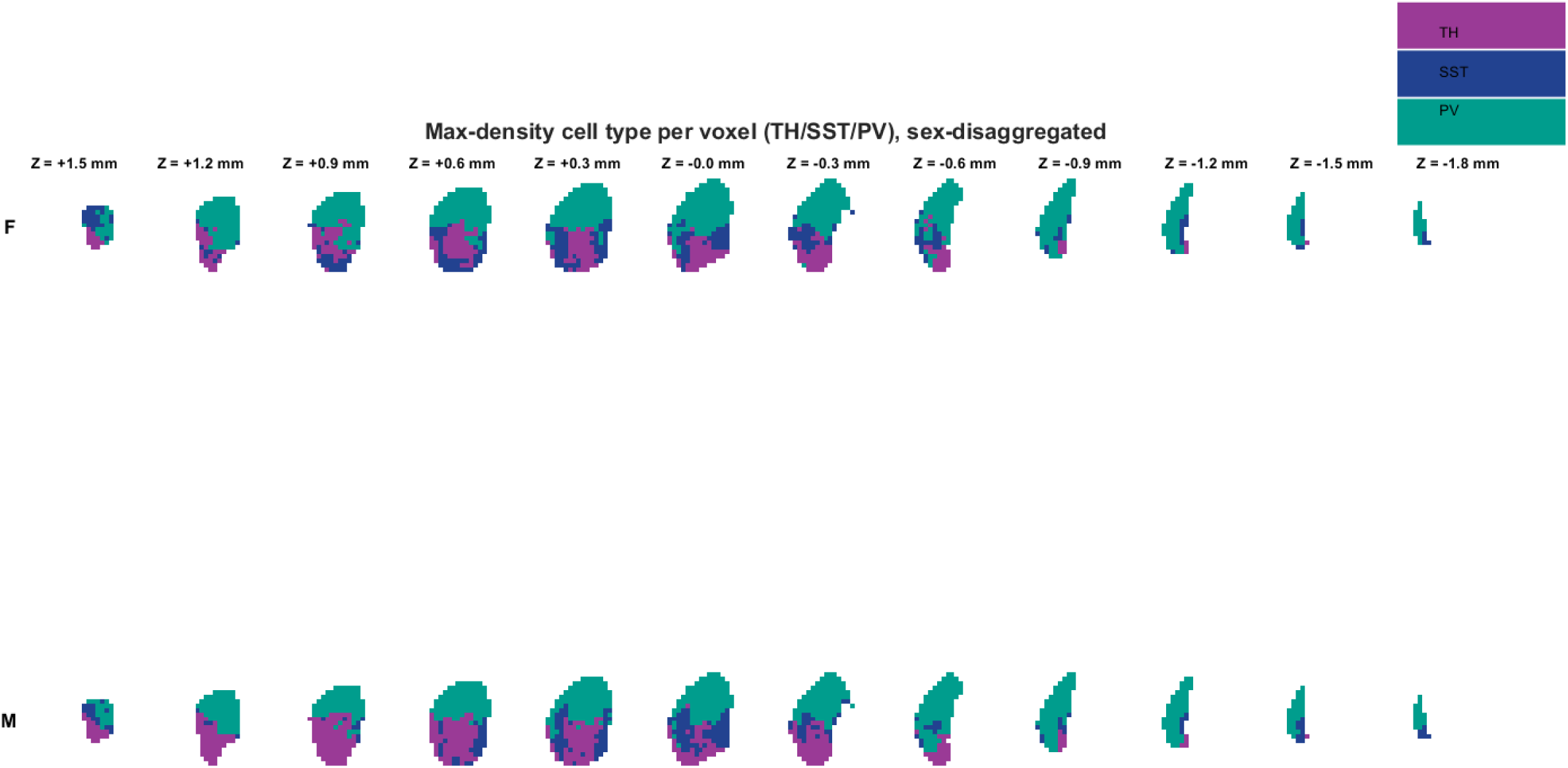
Voxel-wise predominance maps, stratified by sex. Each voxel was assigned to the interneuron subtype exhibiting the highest density. The top row shows voxel-wise interneuron predominance in females (*N*(SST) = 6, *N*(PV) = 5; *N*(TH) = 7), the bottom row in males (*N*(SST) = 6, *N*(PV) = 6; *N*(TH) = 6). Voxels are color-coded to indicate the most abundant interneuron subtype (blue, SST; green, PV; purple, TH). This predominance map provides a qualitative summary of relative enrichment and does not imply exclusivity of interneuron subtypes within individual voxels.

**Supplemental Figure 4.**
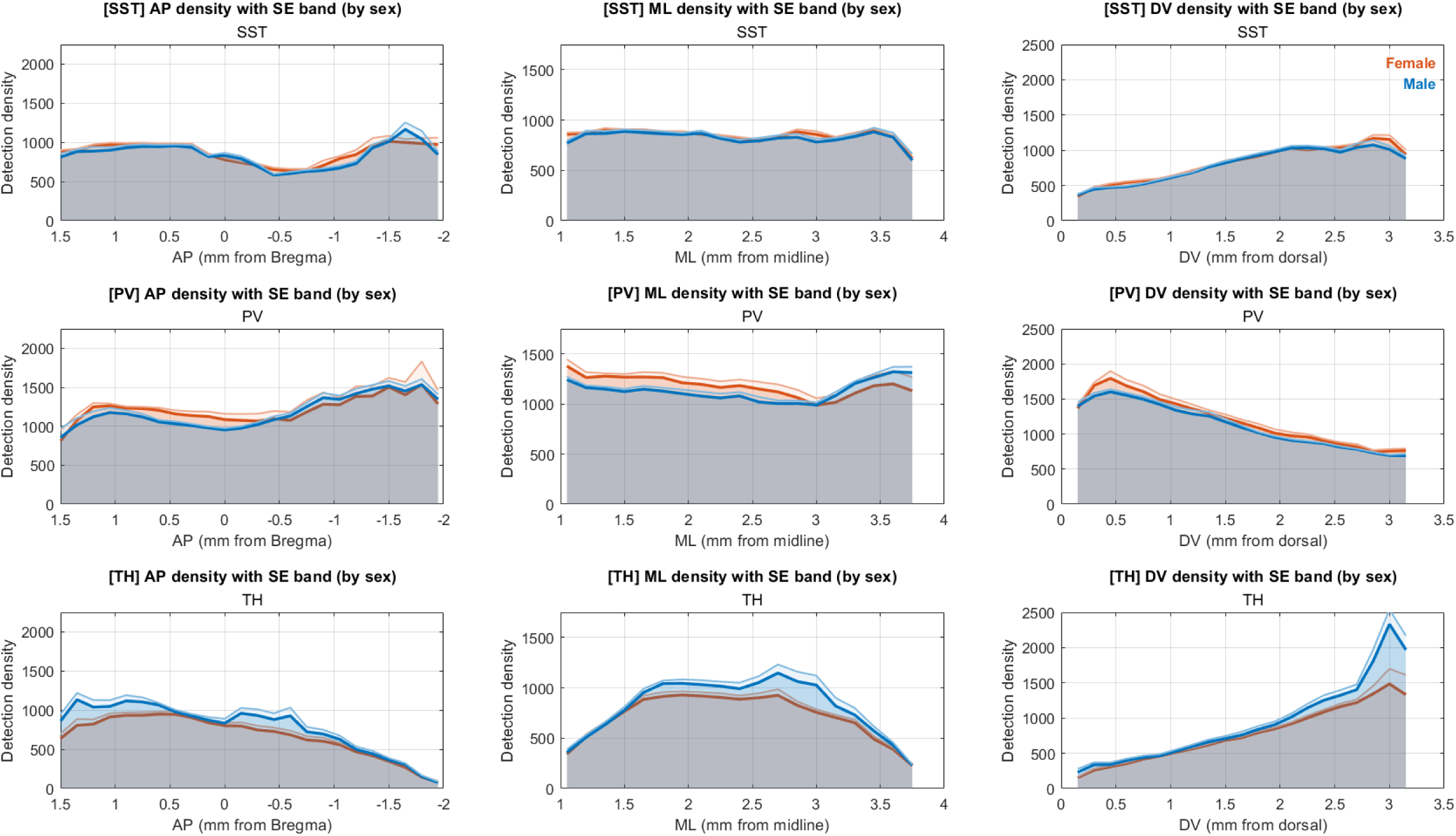
Large-scale distributions of striatal interneuron subtypes across anatomical axes, stratified by sex. Voxel-wise interneuron density estimates were quantified along the anterior-posterior (column 1), medial-lateral (column 2), and dorsal-ventral (column 3) axes of the mouse caudoputamen for SST (row 1), PV (row 2), and TH (row 3) interneurons, in males (blue) and females (orange). Plots depict the bootstrapped mean density across 150-μm planes for SST (*N*(hemispheres) = 12; 6 female, 6 male), PV (*N*(hemispheres) = 11; 5 female, 6 male), and TH (*N*(hemispheres) = 13; 7 female, 6 male) interneurons. Shaded regions indicate mean + s.e. of the bootstrapped mean (1,000 bootstraps per hemisphere). These plots are intended to visualize large-scale spatial trends and facilitate qualitative comparison between sexes; formal statistical analyses of sex effects are reported separately.

**Supplemental Figure 5.**
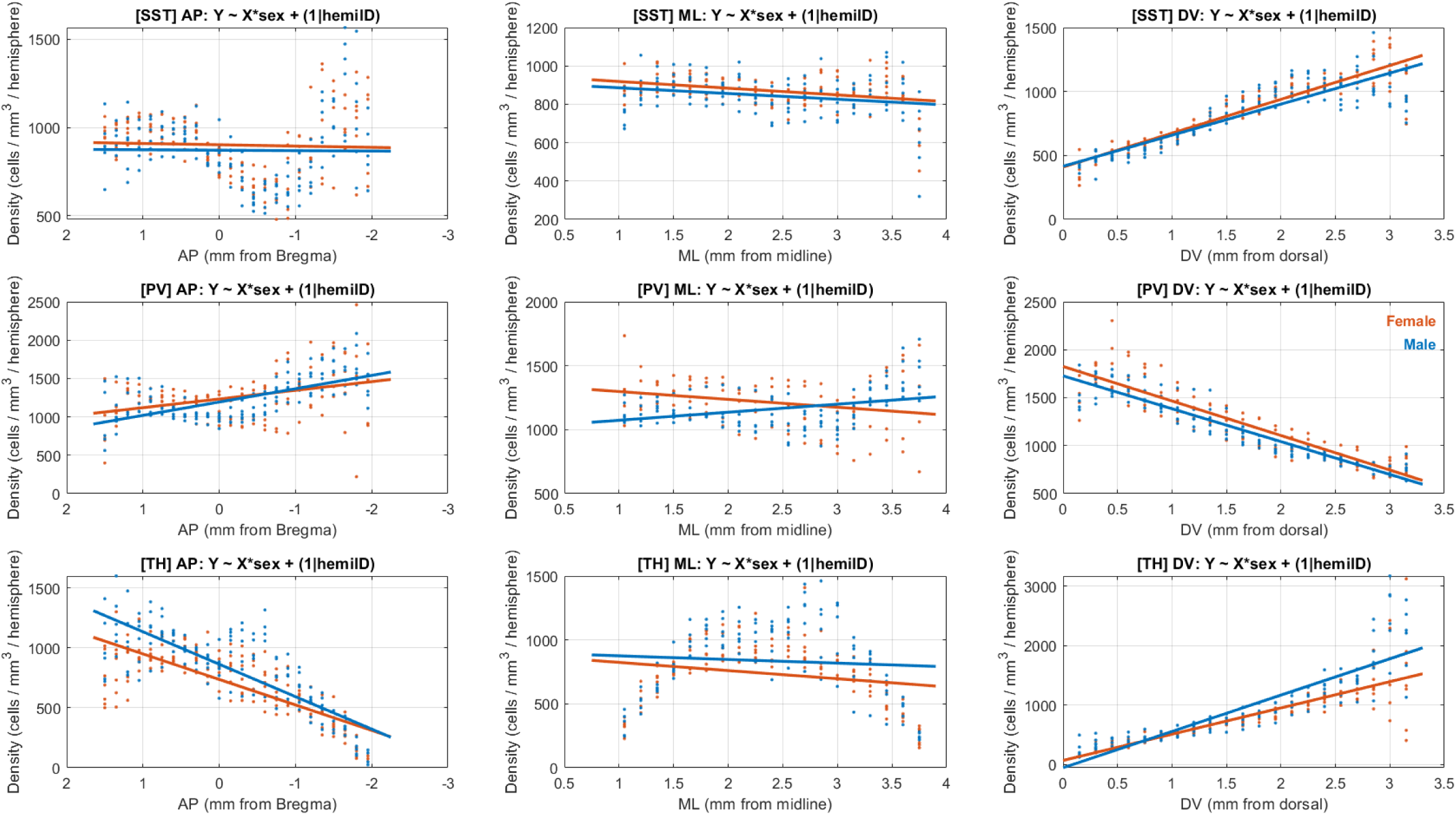
Large-scale spatial gradients of striatal interneuron subtypes across anatomical axes, stratified by sex. Voxel-wise interneuron density estimates were quantified per-hemisphere along the anterior-posterior (column 1), medial-lateral (column 2), and dorsal-ventral (column 3) axes of the mouse caudoputamen for SST (row 1), PV (row 2), and TH (row 3) interneurons, in males (blue) and females (orange). Points indicate per-hemisphere density values for each anatomic plane, with overlaid linear mixed-effects model fits, for SST (*N*(hemispheres) = 12; 6 female, 6 male), PV (*N*(hemispheres) = 11; 5 female, 6 male), and TH (*N*(hemispheres) = 13; 7 female, 6 male) interneurons. Linear mixed effects models (Density ∼ Coordinate * Sex + (1 | Hemisphere_ID)) were used to assess large-scale directional biases while accounting for repeated sampling within hemispheres. Sex-specific slopes were derived from the fitted interaction model, with female slopes corresponding to the Coordinate coefficient, and male slopes corresponding to the sum of the Coordinate and Coordinate x Sex interaction coefficients. Statistical significance of sex-dependent differences in spatial gradients was assessed using the Coordinate x Sum interaction term, with Benjamini-Hochberg false discovery rate (FDR) correction applied across all cell type x axis combinations (q = 0.05). Model slopes represent global directional bias (cells/mm^3^ per mm along each anatomical axis) and are not intended to capture non-monotonic structure. Anterior–posterior axis (AP) SST: Females — β = 7.31 ± 15.60, p = 0.64 (N = 6 hemispheres); Males — β = 2.46 (derived) (N = 6); Sex×AP interaction — Δβ = −4.85 ± 22.06, p = 0.83, q = 0.83. PV: Females — β = −113.10 ± 23.73, p = 3.1 × 10⁻⁶ (N = 5); Males — β = −173.69 (derived) (N = 6); Sex×AP interaction — Δβ = −60.59 ± 32.12, p = 0.060, q = 0.136. TH: Females — β = 212.90 ± 13.03, p = 1.25 × 10⁻⁴³ (N = 7); Males — β = 270.75 (derived) (N = 6); Sex×AP interaction — Δβ = 57.85 ± 19.19, p = 0.00278, q = 0.00834. Medial–lateral axis (ML) SST: Females — β = −35.00 ± 9.93, p = 5.14 × 10⁻⁴ (N = 6); Males — β = −29.78 (derived) (N = 6); Sex×ML interaction — Δβ = 5.22 ± 14.04, p = 0.711, q = 0.800. PV: Females — β = −61.34 ± 19.69, p = 0.00211 (N = 5); Males — β = 62.96 (derived) (N = 6); Sex×ML interaction — Δβ = 124.30 ± 26.67, p = 5.66 × 10⁻⁶, q = 2.55 × 10⁻⁵. TH: Females — β = −63.85 ± 28.89, p = 0.0280 (N = 7); Males — β = −28.40 (derived) (N = 6); Sex×ML interaction — Δβ = 35.45 ± 42.52, p = 0.405, q = 0.521. Dorsal–ventral axis (DV) SST: Females — β = 264.80 ± 10.24, p = 2.16 × 10⁻⁷² (N = 6); Males — β = 243.04 (derived) (N = 6); Sex×DV interaction — Δβ = −21.76 ± 14.48, p = 0.134, q = 0.241. PV: Females — β = −360.30 ± 12.85, p = 1.18 × 10⁻⁷⁵ (N = 5); Males — β = −342.84 (derived) (N = 6); Sex×DV interaction — Δβ = 17.46 ± 17.40, p = 0.317, q = 0.475. TH: Females — β = 441.60 ± 23.38, p = 3.14 × 10⁻⁵¹ (N = 7); Males — β = 610.80 (derived) (N = 6); Sex×DV interaction — Δβ = 169.20 ± 34.42, p = 1.54 × 10⁻⁶, q = 1.39 × 10⁻⁵.

**Supplemental Figure 6.**
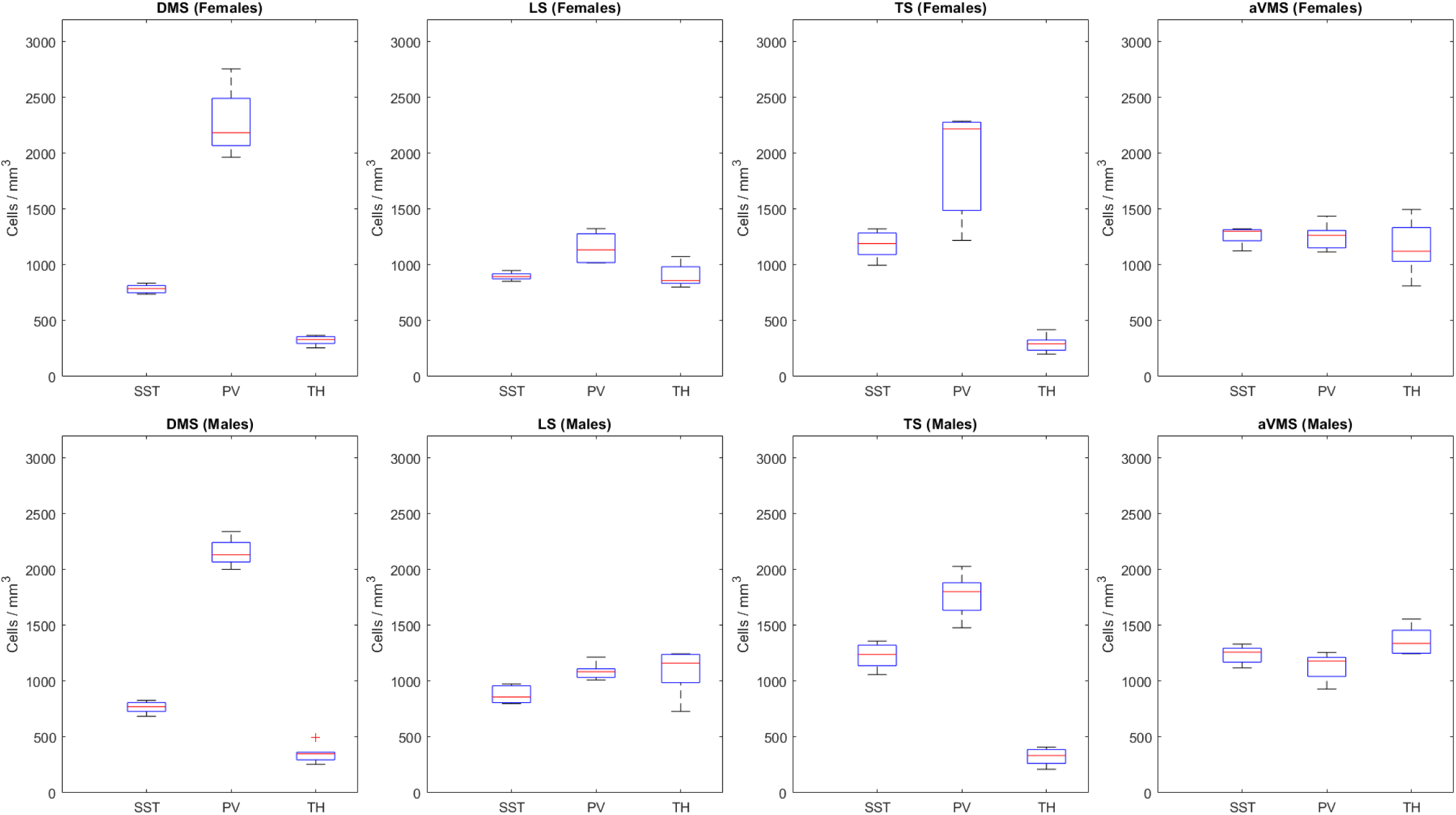
Subregional organization of striatal SST, PV, and TH interneurons, within subregion, stratified by sex. Per-hemisphere subregional density distributions (boxplots; center line, median; box, interquartile range; whiskers, non-outlier extrema; points, outliers) are shown organized by subregion. Females are shown in the top row (*N*(SST) = 6; *N*(PV) = 5; *N*(TH) = 7), males are shown in the bottom row (*N*(SST) = 6; *N*(PV) = 6; *N*(TH) = 6). Columns correspond to dorsomedial striatum (DMS), lateral striatum (LS), tail of striatum (TS), and anterior ventromedial striatum (aVMS), defined using a four-cluster anatomic parcellation (Hunnicutt et al., 2016). Mixed-effects ANOVAs (linear mixed-effects models with random intercept for hemisphere; (1 | Hemisphere_ID)) were used to test sex differences in subtype distributions within each subregion (Density ∼ Subtype * Sex + (1 | Hemisphere_ID)). Significant main effects of interneuron subtype were observed in DMS, DLS, and TS (BH–FDR q < 0.05), but not in aVMS. Significant Subtype × Sex interactions were detected in DLS and aVMS (BH–FDR q < 0.05), indicating sex-dependent modulation of interneuron subtype distributions in these subregions.

**Supplemental Figure 7.**
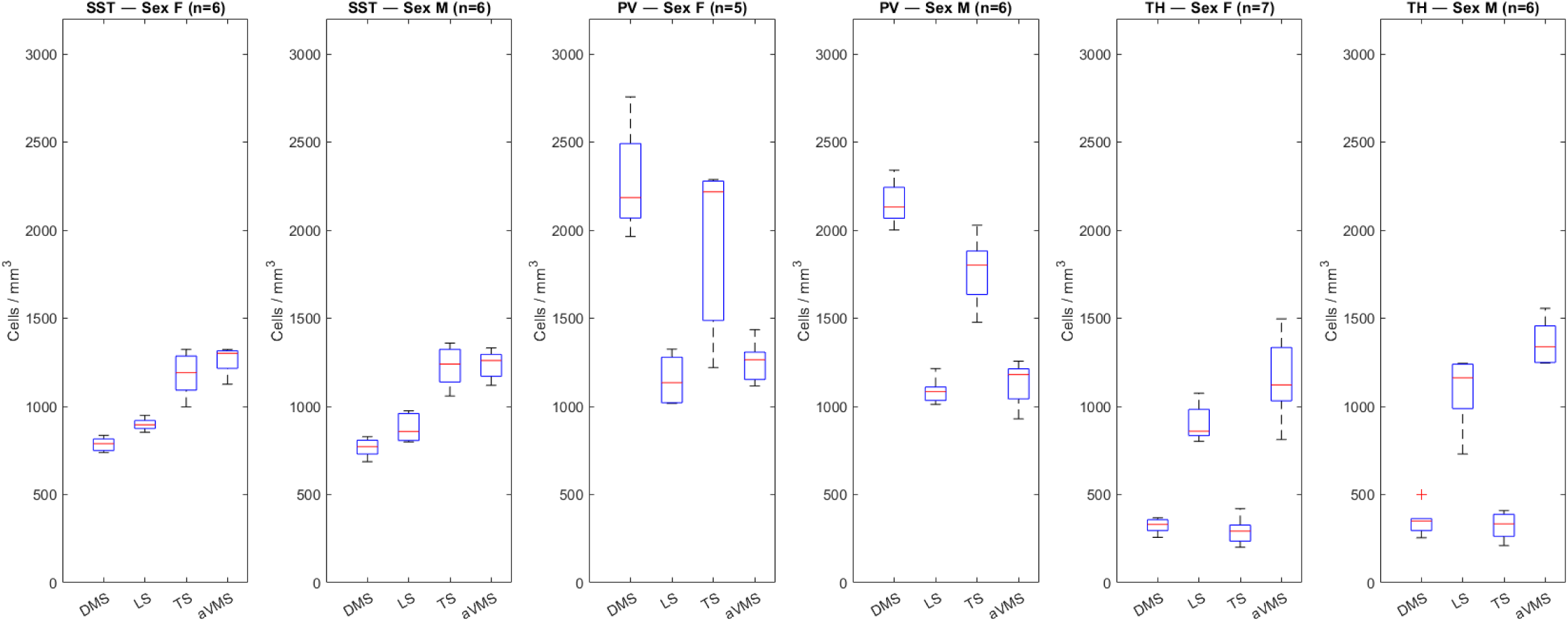
Subregional organization of striatal SST, PV, and TH interneurons, within cell type, stratified by sex. Per-hemisphere subregional density distributions (boxplots; center line, median; box, interquartile range; whiskers, non-outlier extrema; points, outliers) are shown organized by interneuron subtype. Females (*N*(SST) = 6; *N*(PV) = 5; *N*(TH) = 7) and males (*N*(SST) = 6; *N*(PV) = 6; *N*(TH) = 6) are shown in paired box plots for each subtype. Subregions include dorsomedial striatum (DMS), dorsolateral striatum (DLS), tail of striatum (TS), and anterior ventromedial striatum (aVMS), defined using a four-cluster anatomical parcellation (Hunnicutt et al., 2016). Mixed-effects ANOVAs (linear mixed-effects models with random intercept for hemisphere; (1 | Hemisphere_ID)) were used to test sex differences in subregional distribution within each subtype (Density ∼ Subregion * Sex + (1 | Hemisphere_ID)). Significant main effects of subregion were observed for SST, PV, and TH interneurons (BH–FDR q < 0.05), whereas no Subregion × Sex interactions remained significant after FDR correction, indicating conserved subregional organization across sexes.

## Notes

### Competing Interest Statement

The authors have declared no competing interest.

## References

Alexander, G.E., Crutcher, M.D., and DeLong, M.R. (1991). Chapter 6 basal ganglia-thalamocortical circuits: Parallel substrates for motor, oculomotor, “prefrontal” and “limbic” functions. Progress in Brain Research 85. 10.1016/S0079-6123(08)62678-3.

Apicella, P. (2017). The role of the intrinsic cholinergic system of the striatum: What have we learned from tan recordings in behaving animals? Neuroscience 360. 10.1016/j.neuroscience.2017.07.060.

Appings, R., Botterill, J.J., Zhao, M., Riaz, S., Kanani, A., Violi, F., Steininger, C.F.D., Ito, R., and Arruda-Carvalho, M. (2024). Tyrosine hydroxylase–positive nucleus accumbens neurons influence delay discounting in a mouse t-maze task. eNeuro 11. 10.1523/ENEURO.0487-24.2024.

Assous, M., Kaminer, J., Shah, F., Garg, A., Koos, T., and Tepper, J.M. (2017). Differential processing of thalamic information via distinct striatal interneuron circuits. Nat Commun 8, 15860. 10.1038/ncomms15860.

Beckstead, R.M. (1979). An autoradiographic examination of corticocortical and subcortical projections of the mediodorsal-projection (prefrontal) cortex in the rat. Journal of Comparative Neurology 184. 10.1002/cne.901840104.

Bernácer, J., Prensa, L., and Giménez-Amaya, J.M. (2007). Cholinergic interneurons are differentially distributed in the human striatum. PloS one 2. 10.1371/journal.pone.0001174.

Bernácer, J., Prensa, L., and Giménez-Amaya, J.M. (2012). Distribution of gabaergic interneurons and dopaminergic cells in the functional territories of the human striatum. PloS one 7. 10.1371/journal.pone.0030504.

Brimblecombe, K.R., and Cragg, S.J. (2017). The striosome and matrix compartments of the striatum: A path through the labyrinth from neurochemistry toward function. ACS Chem Neurosci 8, 235–242. 10.1021/acschemneuro.6b00333.

Carrasco, A., Oorschot, D.E., Barzaghi, P., and Wickens, J.R. (2022). Three-dimensional spatial analyses of cholinergic neuronal distributions across the mouse septum, nucleus basalis, globus pallidus, nucleus accumbens, and caudate-putamen. Neuroinformatics 20 10.1007/s12021-022-09588-1.

Choi, K., Holly, E., Davatolhagh, M.F., Beier, K.T., and Fuccillo, M.V. (2018). Integrated anatomical and physiological mapping of striatal afferent projections. Eur J Neurosci. 10.1111/ejn.13829.

Choi, K., Piasini, E., Díaz-Hernández, E., Cifuentes, L.V., Henderson, N.T., Holly, E.N., Subramaniyan, M., Gerfen, C.R., and Fuccillo, M.V. (2023). Distributed processing for value-based choice by prelimbic circuits targeting anterior-posterior dorsal striatal subregions in male mice. Nature Communications 14, 1920. 10.1038/s41467-023-36795-4.

Cifuentes, L.V., Díaz-Hernández, E., Granato, M.M., Tu, W., Choi, K., and Fuccillo, M.V. (2025). Temporally and functionally distinct contributions to value based choice along the anterior-posterior dorsomedial striatal axis. bioRxiv. 10.1101/2025.03.14.643367.

Corbit, L.H., and Janak, P.H. (2010). Posterior dorsomedial striatum is critical for both selective instrumental and pavlovian reward learning. Eur J Neurosci 31, 1312–1321. 10.1111/j.1460-9568.2010.07153.x.

Corrigan, E.K., DeBerardine, M., Poddar, A., Turrero García, M., de la O, S., He, S., Sen, H., Duhne, M., Lindberg, S., Song, M., et al. (2025). Conservation and alteration of mammalian striatal interneurons. Nature 647. 10.1038/s41586-025-09592-w.

Cox, J., and Witten, I.B. (2019). Striatal circuits for reward learning and decision-making. Nature reviews. Neuroscience 20, 482–494. 10.1038/s41583-019-0189-2.

Duhne, M., Lara-González, E., Laville, A., Padilla-Orozco, M., Ávila-Cascajares, F., Arias-García, M., Galarraga, E., and Bargas, J. (2021). Activation of parvalbumin-expressing neurons reconfigures neuronal ensembles in murine striatal microcircuits. European Journal of Neuroscience 53. 10.1111/ejn.14670.

Duhne, M., Mohebi, A., Kim, K., Pelattini, L., and Berke, J.D. (2024). A mismatch between striatal cholinergic pauses and dopaminergic reward prediction errors. Proceedings of the National Academy of Sciences 121. 10.1073/pnas.2410828121.

Duhne, M., Montalvo, I.G., Zheng, C., Pelattini, L., and Berke, J.D. (2025). Parvalbumin interneurons gate and shape striatal sequences. bioRxiv. 10.1101/2025.11.08.687375.

Eastwood, B.S., Hooks, B.M., Paletzki, R.F., O’Connor, N.J., Glaser, J.R., and Gerfen, C.R. (2018). Whole mouse brain reconstruction and registration to a reference atlas with standard histochemical processing of coronal sections. The Journal of comparative neurology. 10.1002/cne.24602.

Fino, E., Vandecasteele, M., Perez, S., Saudou, F., and Venance, L. (2018). Region-specific and state-dependent action of striatal gabaergic interneurons. Nat Commun 9, 3339. 10.1038/s41467-018-05847-5.

Garas, F.N., Shah, R.S., Kormann, E., Doig, N.M., Vinciati, F., Nakamura, K.C., Dorst, M.C., Smith, Y., Magill, P.J., and Sharott, A. (2016). Secretagogin expression delineates functionally-specialized populations of striatal parvalbumin-containing interneurons. eLife 5. 10.7554/eLife.16088.

Gerfen, C.R. (1985). The neostriatal mosaic. I. Compartmental organization of projections from the striatum to the substantia nigra in the rat. Journal of Comparative Neurology 236. 10.1002/cne.902360404.

Graybiel, A.M., and Grafton, S.T. (2015). The striatum: Where skills and habits meet. Cold Spring Harbor perspectives in biology 7. 10.1101/cshperspect.a021691.

Graybiel, A.M., and Matsushima, A. (2023). Striosomes and matrisomes: Scaffolds for dynamic coupling of volition and action. Annual Review of Neuroscience 46. 10.1146/annurev-neuro-121522-025740.

Gritton, H.J., Howe, W.M., Romano, M.F., Difeliceantonio, A.G., Kramer, M.A., Saligrama, V., Bucklin, M.E., Zemel, D., and Han, X. (2019). Unique contributions of parvalbumin and cholinergic interneurons in organizing striatal networks during movement. Nature neuroscience 22, 586–597. 10.1038/s41593-019-0341-3.

Hintiryan, H., Foster, N.N., Bowman, I., Bay, M., Song, M.Y., Gou, L., Yamashita, S., Bienkowski, M.S., Zingg, B., Zhu, M., et al. (2016). The mouse cortico-striatal projectome. Nature neuroscience 19, 1100–1114. 10.1038/nn.4332.

Hobel, Z.B., Yang, L.-T., Brechbill, T.R., Liu, Q., Goldberg, J.A., and Plotkin, J.L. (2026). Striatal cholinergic interneurons exhibit compartment-specific anatomical and functional organization in the mouse. Proceedings of the National Academy of Sciences 123. 10.1073/pnas.2519939123.

Holly, E.N., Davatolhagh, M.F., Choi, K., Alabi, O.O., Vargas Cifuentes, L., and Fuccillo, M.V. (2019). Striatal low-threshold spiking interneurons regulate goal-directed learning. Neuron. 10.1016/j.neuron.2019.04.016.

Holly, E.N., Davatolhagh, M.F., Espana, R.A., and Fuccillo, M.V. (2021). Striatal low-threshold spiking interneurons locally gate dopamine. Curr Biol. 10.1016/j.cub.2021.06.081.

Hunnicutt, B.J., Jongbloets, B.C., Birdsong, W.T., Gertz, K.J., Zhong, H., and Mao, T. (2016). A comprehensive excitatory input map of the striatum reveals novel functional organization. Elife 5. 10.7554/eLife.19103.

Ibanez-Sandoval, O., Tecuapetla, F., Unal, B., Shah, F., Koos, T., and Tepper, J.M. (2010). Electrophysiological and morphological characteristics and synaptic connectivity of tyrosine hydroxylase-expressing neurons in adult mouse striatum. Journal of Neuroscience 30, 6999–7016.

Kaminer, J., Espinoza, D., Bhimani, S., Tepper, J.M., Koos, T., and Shiflett, M.W. (2019). Loss of striatal tyrosine-hydroxylase interneurons impairs instrumental goal-directed behavior. European Journal of Neuroscience 50, 2653–2662. 10.1111/ejn.14412.

Klug, J.R., Engelhardt, M.D., Cadman, C.N., Li, H., Smith, J.B., Ayala, S., Williams, E.W., Hoffman, H., and Jin, X. (2018). Differential inputs to striatal cholinergic and parvalbumin interneurons imply functional distinctions. eLife 7. 10.7554/eLife.35657.

Kobayashi, Y., and Hensch, T.K. (2013). Germline recombination by conditional gene targeting with parvalbumin-cre lines. Frontiers in Neural Circuits 7. 10.3389/fncir.2013.00168.

Kruttner, S., Falasconi, A., Valbuena, S., Galimberti, I., Bouwmeester, T., Arber, S., and Caroni, P. (2022). Absence of familiarity triggers hallmarks of autism in mouse model through aberrant tail-of-striatum and prelimbic cortex signaling. Neuron 110, 1468–1482 e1465. 10.1016/j.neuron.2022.02.001.

Lecumberri, A., Lopez-Janeiro, A., Corral-Domenge, C., and Bernacer, J. (2017). Neuronal density and proportion of interneurons in the associative, sensorimotor and limbic human striatum. Neuroscience 223. 10.1007/s00429-017-1579-8.

Lee, J., and Sabatini, B.L. (2025). From avoidance to new action: The multifaceted role of the striatal indirect pathway. Nature Reviews Neuroscience 26. 10.1038/s41583-025-00925-2.

Lee, K., Holley, S.M., Shobe, J.L., Chong, N.C., Cepeda, C., Levine, M.S., and Masmanidis, S.C. (2017). Parvalbumin interneurons modulate striatal output and enhance performance during associative learning. Neuron 93, 1451–1463 e1454. 10.1016/j.neuron.2017.02.033.

Lein, E.S., Hawrylycz, M.J., Ao, N., Ayres, M., Bensinger, A., Bernard, A., Boe, A.F., Boguski, M.S., Brockway, K.S., Byrnes, E.J., et al. (2006). Genome-wide atlas of gene expression in the adult mouse brain. Nature 445. 10.1038/nature05453.

López-González del Rey, N., Trigo-Damas, I., Obeso, J.A., Cavada, C., and Blesa, J. (2022). Neuron types in the primate striatum: Stereological analysis of projection neurons and interneurons in control and parkinsonian monkeys. Neuropathology and Applied Neurobiology 48. 10.1111/nan.12812.

Luk, K.C., and Sadikot, A.F. (2001). Gaba promotes survival but not proliferation of parvalbumin-immunoreactive interneurons in rodent neostriatum: An in vivo study with stereology. Neuroscience 104, 93–103.

Martel, A.-C., and Galvan, A. (2022). Connectivity of the corticostriatal and thalamostriatal systems in normal and parkinsonian states: An update. Neurobiology of Disease 174. 10.1016/j.nbd.2022.105878.

Martiros, N., Burgess, A.A., and Graybiel, A.M. (2018). Inversely active striatal projection neurons and interneurons selectively delimit useful behavioral sequences. Current Biology 28. 10.1016/j.cub.2018.01.031.

Matamales, M., Götz, J., and Bertran-Gonzalez, J. (2016). Quantitative imaging of cholinergic interneurons reveals a distinctive spatial organization and a functional gradient across the mouse striatum. PloS one 11. 10.1371/journal.pone.0157682.

Mestres-Missé, A., Turner, R., and Friederici, A.D. (2012). An anterior–posterior gradient of cognitive control within the dorsomedial striatum. NeuroImage 62. 10.1016/j.neuroimage.2012.05.021.

Monteiro, P., Barak, B., Zhou, Y., McRae, R., Rodrigues, D., Wickersham, I.R., and Feng, G. (2018). Dichotomous parvalbumin interneuron populations in dorsolateral and dorsomedial striatum. The Journal of Physiology 596. 10.1113/JP275936.

Muñoz-Manchado, A.B., Bengtsson Gonzales, C., Zeisel, A., Munguba, H., Bekkouche, B., Skene, N.G., Lönnerberg, P., Ryge, J., Harris, K.D., Linnarsson, S., and Hjerling-Leffler, J. (2018). Diversity of interneurons in the dorsal striatum revealed by single-cell rna sequencing and patchseq. Cell Reports 24, 2179–2190.e2177. 10.1016/j.celrep.2018.07.053.

O’Hare, J.K., Li, H., Kim, N., Gaidis, E., Ade, K., Beck, J., Yin, H., and Calakos, N. (2017). Striatal fast-spiking interneurons selectively modulate circuit output and are required for habitual behavior. Elife 6. 10.7554/eLife.26231.

Owen, S.F., Berke, J.D., and Kreitzer, A.C. (2018). Fast-spiking interneurons supply feedforward control of bursting, calcium, and plasticity for efficient learning. Cell 172, 683–695 e615. 10.1016/j.cell.2018.01.005.

Rotariu, S., Zalcman, G., Badreddine, N., Appaix, F., Sarno, S., Bureau, I., and Fino, E. (2025). Somatostatin interneurons select dorsomedial striatal representations of the initial motor learning phase. Cell Reports 44. 10.1016/j.celrep.2025.115670.

Selemon, L.D., and Goldman-Rakic, P.S. (1985). Longitudinal topography and interdigitation of corticostriatal projections in the rhesus monkey. The Journal of neuroscience: the official journal of the Society for Neuroscience 5, 776–794.

Shiflett, M.W., Brown, R.A., and Balleine, B.W. (2010). Acquisition and performance of goal-directed instrumental actions depends on erk signaling in distinct regions of dorsal striatum in rats. Journal of Neuroscience 30. 10.1523/JNEUROSCI.1778-09.2010.

Straub, C., Saulnier, J.L., Begue, A., Feng, D.D., Huang, K.W., and Sabatini, B.L. (2016). Principles of synaptic organization of gabaergic interneurons in the striatum. Neuron 92, 84–92. 10.1016/j.neuron.2016.09.007.

Tepper, J.M., Koos, T., Ibanez-Sandoval, O., Tecuapetla, F., Faust, T.W., and Assous, M. (2018). Heterogeneity and diversity of striatal gabaergic interneurons: Update 2018. Front Neuroanat 12, 91. 10.3389/fnana.2018.00091.

Tepper, J.M., Tecuapetla, F., Koos, T., and Ibanez-Sandoval, O. (2010). Heterogeneity and diversity of striatal gabaergic interneurons. Front Neuroanat 4.

Unal, B., Ibáñez-Sandoval, O., Shah, F., Abercrombie, E., and Tepper, J.M. (2011). Distribution of tyrosine hydroxylase-expressing interneurons with respect to anatomical organization of the neostriatum. Frontiers in Systems Neuroscience 5. 10.3389/fnsys.2011.00041.

Van Zandt, M., Flanagan, D., and Pittenger, C. (2024). Sex differences in the distribution and density of regulatory interneurons in the striatum. Frontiers in Cellular Neuroscience 18. 10.3389/fncel.2024.1415015.

Wall, N.R., De La Parra, M., Callaway, E.M., and Kreitzer, A.C. (2013). Differential innervation of direct- and indirect-pathway striatal projection neurons. Neuron 79, 347–360.

Wang, X., Qiao, Y., Dai, Z., Sui, N., Shen, F., Zhang, J., and Liang, J. (2018). Medium spiny neurons of the anterior dorsomedial striatum mediate reversal learning in a cell-type-dependent manner. Brain Structure and Function 224. 10.1007/s00429-018-1780-4.

Wu, Y., and Parent, A. (2000). Striatal interneurons expressing calretinin, parvalbumin or nadph-diaphorase: A comparative study in the rat, monkey and human. Brain Research 863. 10.1016/S0006-8993(00)02135-1.

Xenias, H.S., Ibáñez-Sandoval, O., Koós, T., and Tepper, J.M. (2015). Are striatal tyrosine hydroxylase interneurons dopaminergic? The Journal of Neuroscience 35, 6584–6599. 10.1523/jneurosci.0195-15.2015.

Yin, H.H., Ostlund, S.B., Knowlton, B.J., and Balleine, B.W. (2005). The role of the dorsomedial striatum in instrumental conditioning. Eur J Neurosci 22, 513–523.

