## Supplement for "Comprehensive 3D mapping reveals distinct spatial gradients of SST, PV, and TH interneurons across the mouse caudoputamen"

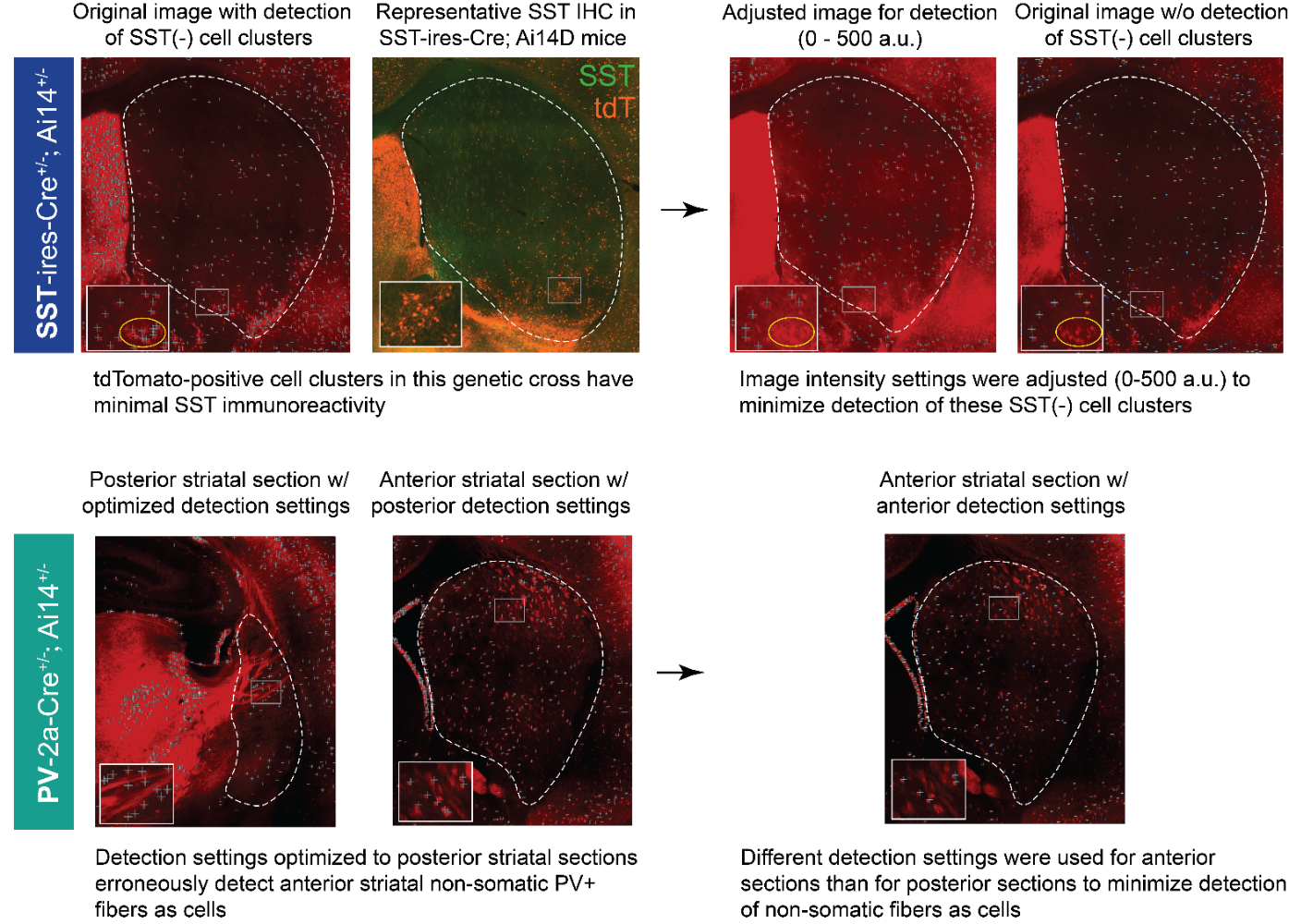

**Supplemental Figure 1. Optimization of detection parameters for SST and PV interneurons.**

Row 1 (SST interneurons): In SST-ires-Cre; Ai14D mice, tdTomato-positive cell clusters were observed that lacked detectable SST immunoreactivity. To restrict automated detection to SST-immunopositive somata, image intensity display ranges were adjusted (0-500 a.u.) such that SST-negative clusters were excluded from detection. This approach substantially reduced their contributions to final SST interneuron counts.

Row 2 (PV interneurons): Non-somatic PV+ fiber bundles exhibit distinct morphology and vary along the anteroposterior axis of the striatum. Detection parameters optimized for posterior sections minimized inclusion of these fibers but resulted in over-detection when applied to anterior planes. To avoid systemic inclusion of non-somatic PV-positive structures, separate detection sensitivity settings were applied for anterior, middle, and posterior sections, ensuring preferential detection of PV-positive somata across the full anteroposterior extent of the striatum.

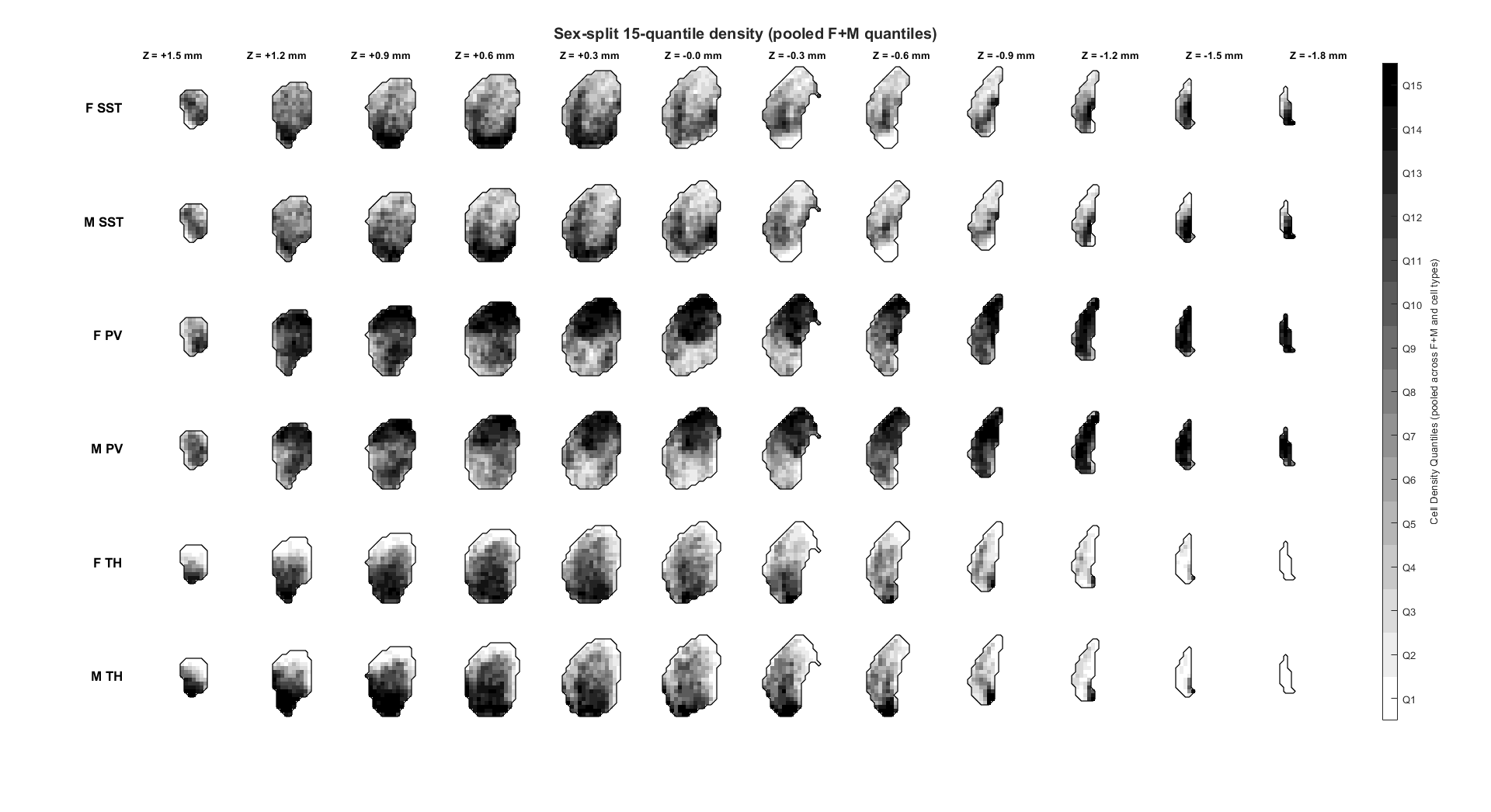
**Supplemental Figure 2. Comprehensive three-dimensional atlas of SST, PV, and TH interneuron distribution across the mouse caudoputamen, stratified by sex.**

Voxel-wise density maps showing the spatial distribution of SST (*N*(hemispheres) = 12; 6 female, 6 male), PV (*N*(hemispheres) = 11; 5 female, 6 male), and TH (*N*(hemispheres) = 13; 7 female, 6 male) interneurons across the mouse caudoputamen. The caudoputamen was partitioned into 150-μm voxels, and interneuron densities were computed for each voxel. For visualization purposes, density values were pooled across cell types and sexes to generate 15 quantile-based density thresholds, displayed from sparse to dense. This common quantile scale enables qualitative comparison of large-scale spatial organization across interneuron subtypes and sexes.

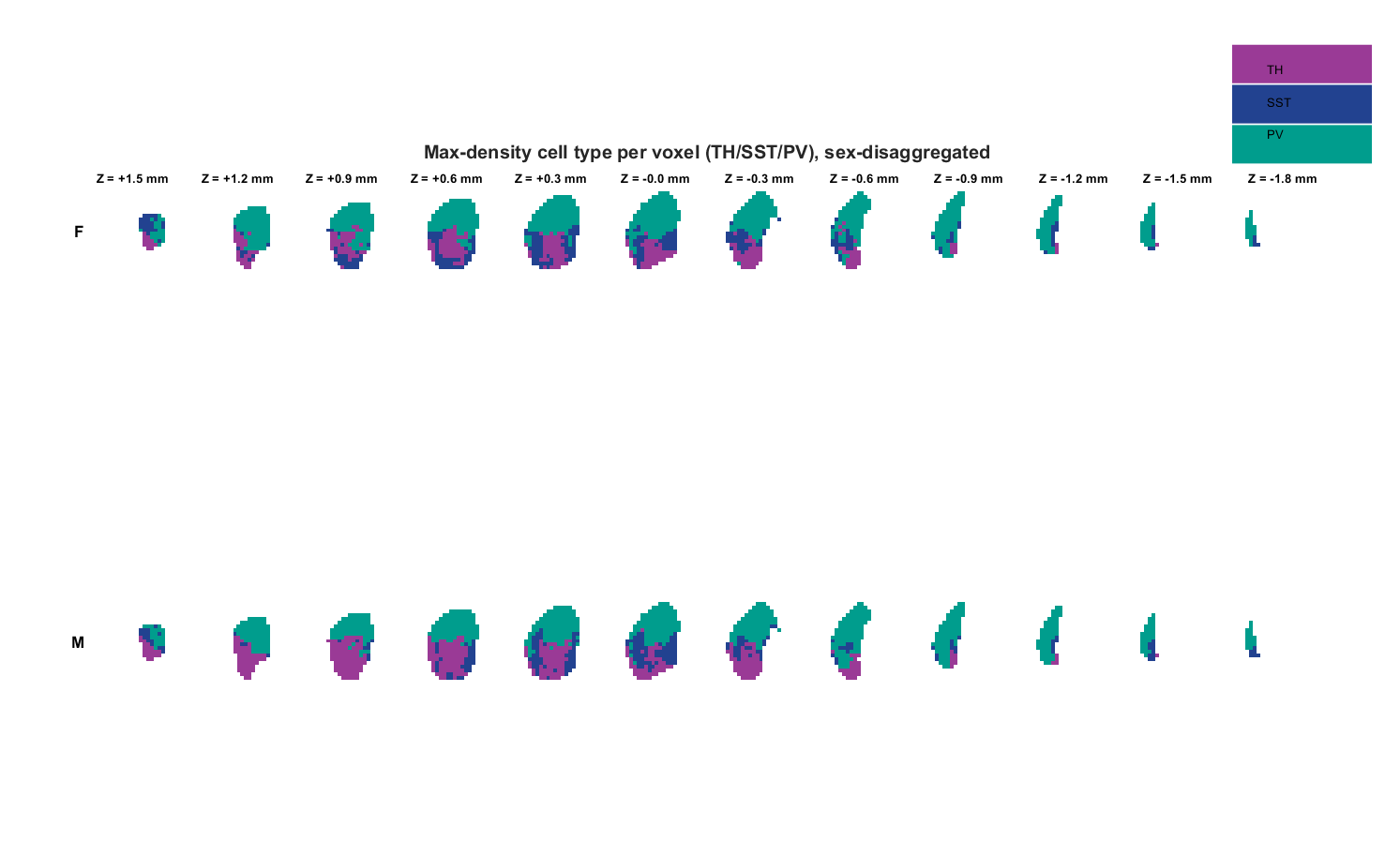

**Supplemental Figure 3. Voxel-wise predominance maps, stratified by sex.**

Each voxel was assigned to the interneuron subtype exhibiting the highest density. The top row shows voxel-wise interneuron predominance in females (*N*(SST) = 6, *N*(PV) = 5; *N*(TH) = 7), the bottom row in males (*N*(SST) = 6, *N*(PV) = 6; *N*(TH) = 6). Voxels are color-coded to indicate the most abundant interneuron subtype (blue, SST; green, PV; purple, TH). This predominance map provides a qualitative summary of relative enrichment and does not imply exclusivity of interneuron subtypes within individual voxels.

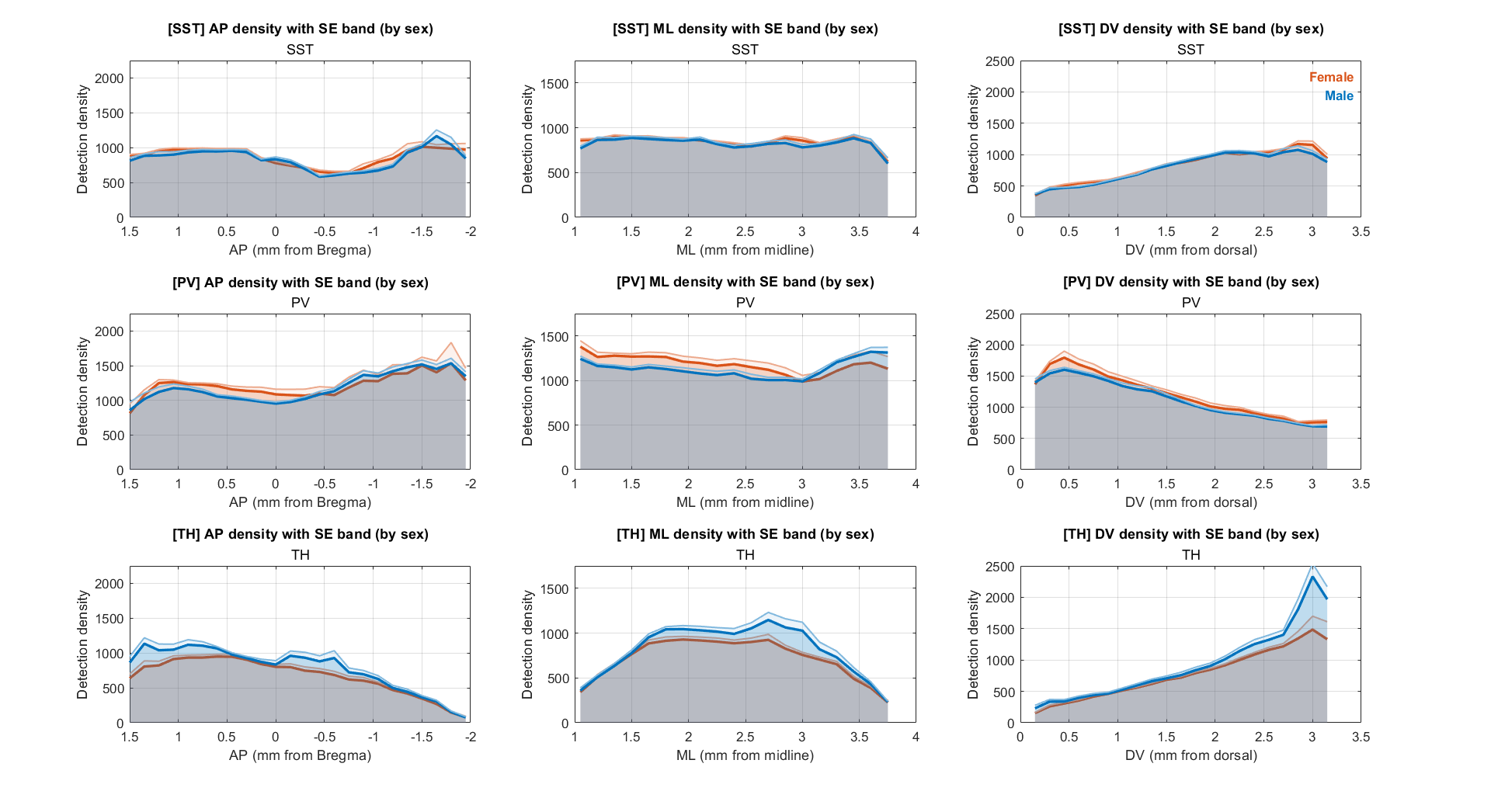

**Supplemental Figure 4. Large-scale distributions of striatal interneuron subtypes across anatomical axes, stratified by sex.**

Voxel-wise interneuron density estimates were quantified along the anterior-posterior (column 1), medial-lateral (column 2), and dorsal-ventral (column 3) axes of the mouse caudoputamen for SST (row 1), PV (row 2), and TH (row 3) interneurons, in males (blue) and females (orange). Plots depict the bootstrapped mean density across 150-μm planes for SST (*N*(hemispheres) = 12; 6 female, 6 male), PV (*N*(hemispheres) = 11; 5 female, 6 male), and TH (*N*(hemispheres) = 13; 7 female, 6 male) interneurons. Shaded regions indicate mean + s.e. of the bootstrapped mean (1,000 bootstraps per hemisphere). These plots are intended to visualize large-scale spatial trends and facilitate qualitative comparison between sexes; formal statistical analyses of sex effects are reported separately.

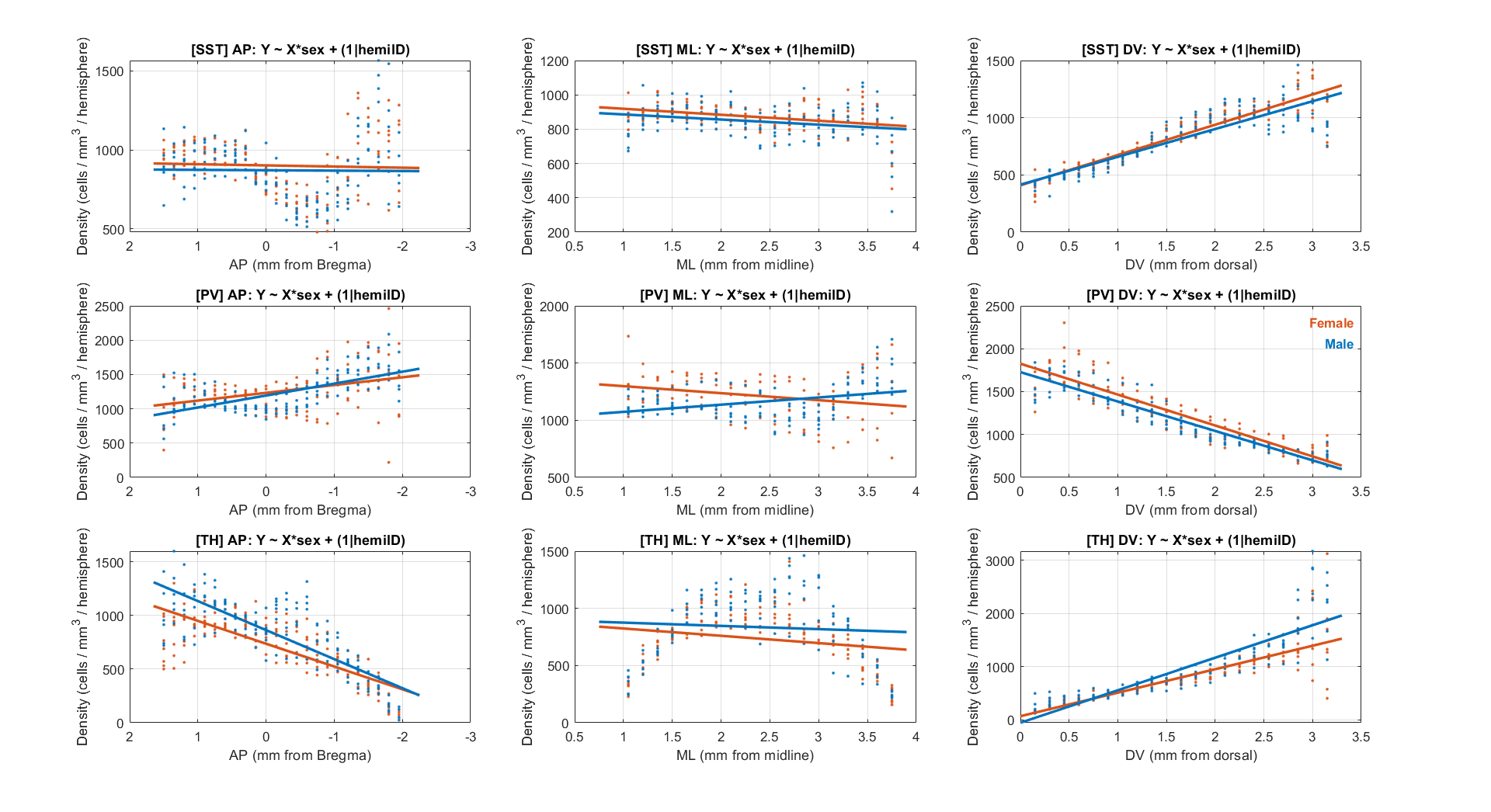
**Supplemental Figure 5. Large-scale spatial gradients of striatal interneuron subtypes across anatomical axes, stratified by sex.**

Voxel-wise interneuron density estimates were quantified per-hemisphere along the anterior-posterior (column 1), medial-lateral (column 2), and dorsal-ventral (column 3) axes of the mouse caudoputamen for SST (row 1), PV (row 2), and TH (row 3) interneurons, in males (blue) and females (orange). Points indicate per-hemisphere density values for each anatomic plane, with overlaid linear mixed-effects model fits, for SST (*N*(hemispheres) = 12; 6 female, 6 male), PV (*N*(hemispheres) = 11; 5 female, 6 male), and TH (*N*(hemispheres) = 13; 7 female, 6 male) interneurons.

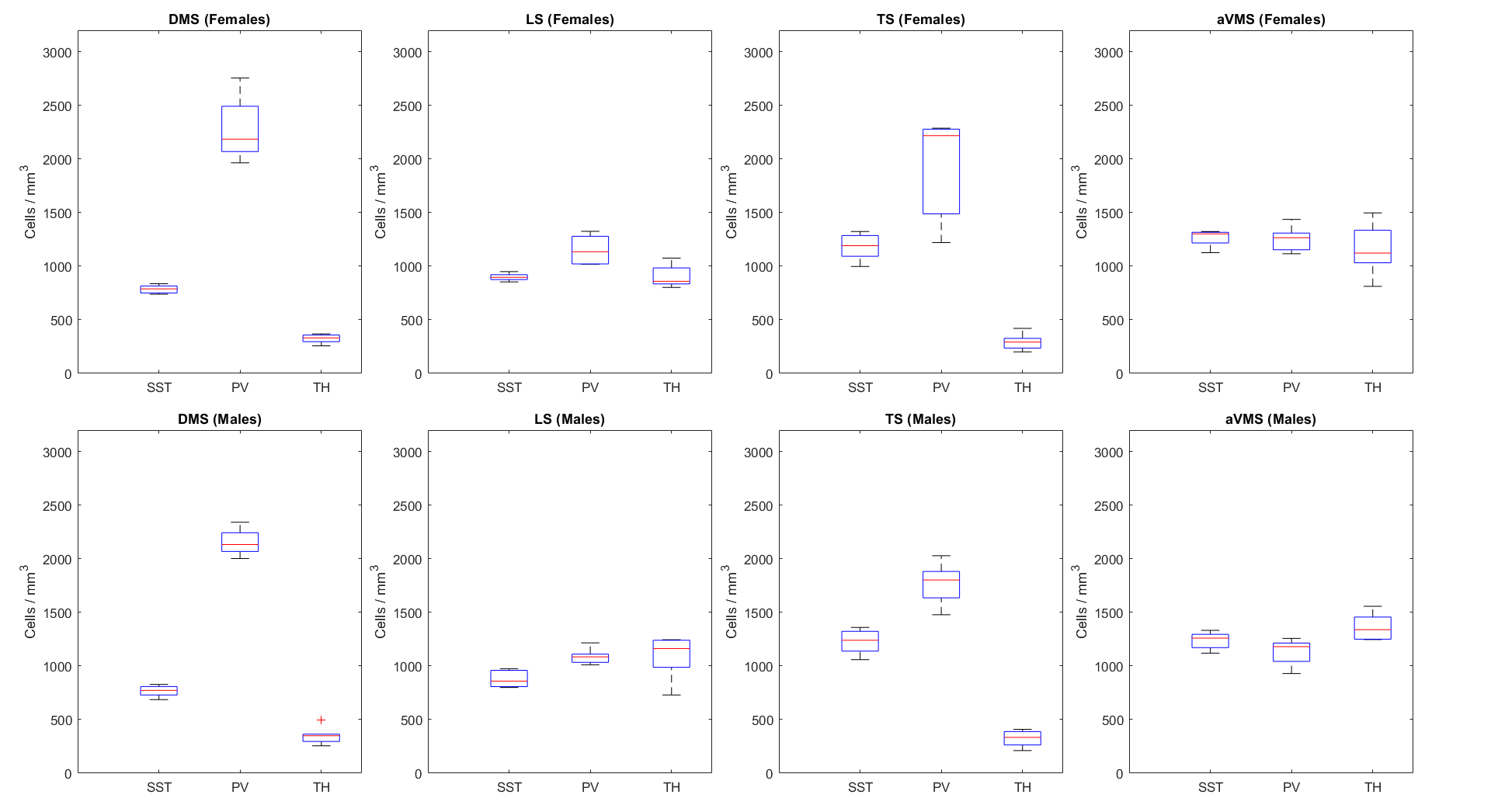

**Supplemental Figure 6. Subregional organization of striatal SST, PV, and TH interneurons, within subregion, stratified by sex.** Per-hemisphere subregional density distributions (boxplots; center line, median; box, interquartile range; whiskers, non-outlier extrema; points, outliers) are shown organized by subregion. Females are shown in the top row (*N*(SST) = 6; *N*(PV) = 5; *N*(TH) = 7), males are shown in the bottom row (*N*(SST) = 6; *N*(PV) = 6; *N*(TH) = 6). Columns correspond to dorsomedial striatum (DMS), lateral striatum (LS), tail of striatum (TS), and anterior ventromedial striatum (aVMS), defined using a four-cluster anatomic parcellation (Hunnicutt et al., 2016). Mixed-effects ANOVAs (linear mixed-effects models with random intercept for hemisphere; (1 | Hemisphere_ID)) were used to test sex differences in subtype distributions within each subregion (Density ~ Subtype * Sex + (1 | Hemisphere_ID)). Significant main effects of interneuron subtype were observed in DMS, DLS, and TS (BH–FDR q < 0.05), but not in aVMS. Significant Subtype × Sex interactions were detected in DLS and aVMS (BH–FDR q < 0.05), indicating sex-dependent modulation of interneuron subtype distributions in these subregions.

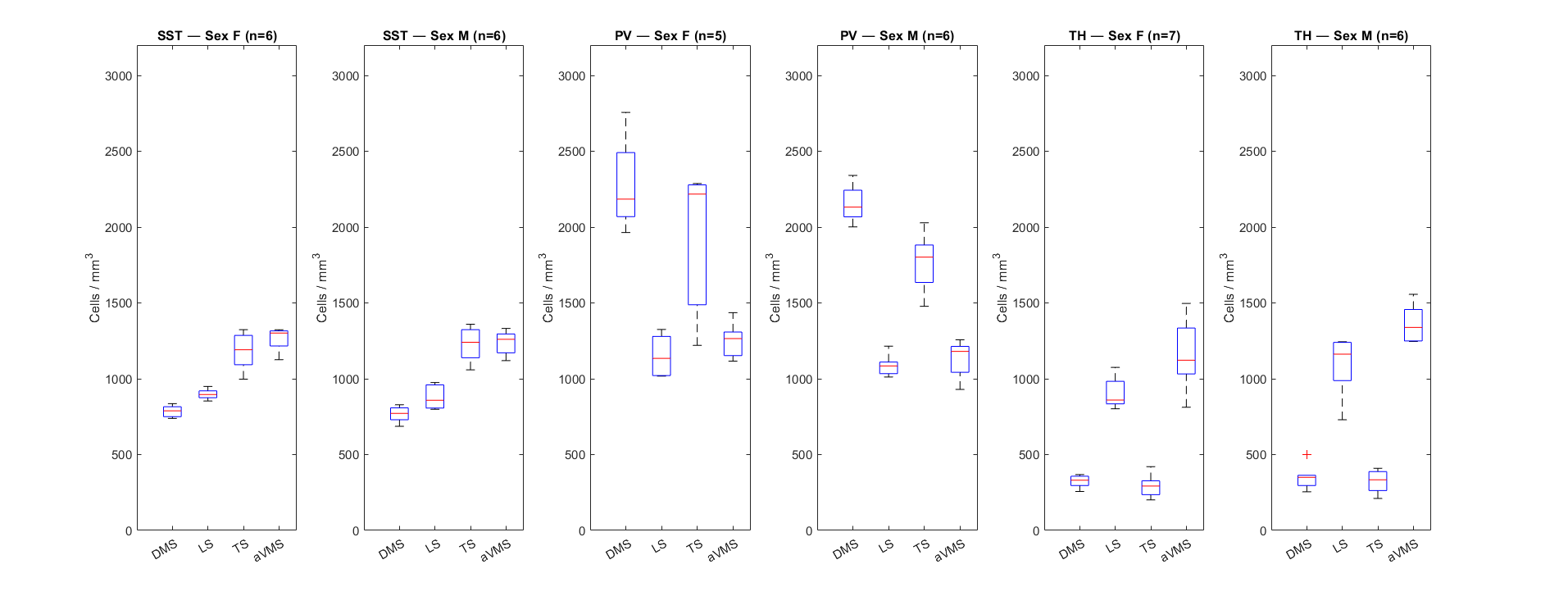

**Supplemental Figure 7. Subregional organization of striatal SST, PV, and TH interneurons, within cell type, stratified by sex**.
Per-hemisphere subregional density distributions (boxplots; center line, median; box, interquartile range; whiskers, non-outlier extrema; points, outliers) are shown organized by interneuron subtype. Females (*N*(SST) = 6; *N*(PV) = 5; *N*(TH) = 7) and males (*N*(SST) = 6; *N*(PV) = 6; *N*(TH) = 6) are shown in paired box plots for each subtype. Subregions include dorsomedial striatum (DMS), dorsolateral striatum (DLS), tail of striatum (TS), and anterior ventromedial striatum (aVMS), defined using a four-cluster anatomical parcellation (Hunnicutt et al., 2016). Mixed-effects ANOVAs (linear mixed-effects models with random intercept for hemisphere; (1 | Hemisphere_ID)) were used to test sex differences in subregional distribution within each subtype (Density ~ Subregion * Sex + (1 | Hemisphere_ID)). Significant main effects of subregion were observed for SST, PV, and TH interneurons (BH–FDR q < 0.05), whereas no Subregion × Sex interactions remained significant after FDR correction, indicating conserved subregional organization across sexes.
